# Cell polarity control by an unconventional G-protein complex in bacteria

**DOI:** 10.1101/2024.10.31.621274

**Authors:** Céline Dinet, Corinne Sebban-Kreuzer, Deborah Byrne-Kodjabachian, Sébastien Lhospice, Julien Herrou, Fabien Durbesson, Renaud Vincentelli, Alphée Michelot, Tâm Mignot

## Abstract

In *Myxococcus xanthus*, cell polarity and motility are regulated by the small GTPase MglA and its associated regulators, MglB, a potential GTPase Activating Protein and the RomRX complex, a potential Guanine nucleotide Exchange Factor. However, recent results have questioned the exact function of these regulators. Using a new type of fluorescent nucleotides for fluorescence-based anisotropy, we first demonstrate that RomRX does not function as a GEF but as an effector that binds strictly to MglA_GTP_. Secondly, based on an enzymology model, we found that MglB functions as a weak catalyst, unlike typical GAPs, and requires additional factors to dissociate from MglA. We demonstrate that RomRX, and specifically RomR, effectively competes with MglA by sequestering MglB. This low affinity interaction is still permissive for GAP activity but sufficient to partition these proteins *in vivo* as indicated by a biomimetic assay in oil emulsion droplets reconstituting these interactions in a cell-sized environment. These results suggest a new model for cell polarity where RomRX exerts a dual function, targeting MglA_GTP_ to the pole via its effector function and preventing its accumulation at the lagging pole by destabilizing the MglA-MglB complex.

## Introduction

Cell polarity, the asymmetric organization of cellular components, is a fundamental characteristic of many living organisms. It is essential for effective cell motility and a myriad of other directional functions, such as epithelial transport, neuronal signaling, and body axis specification in embryos (Allam *et al*, 2018; Mayor & Etienne-Manneville, 2016). A polarized cell establishes a distinct axis with a specific front, known as the polarity site, enabling it to maintain functionally distinct domains. This ability relies on intricate interactions between various protein complexes, including small GTPases, polarity proteins, and cytoskeletal components, that regulate multiple signaling pathways. In most eukaryotic cells, small GTPases of the Ras superfamily act as molecular switches in many signaling pathways and play a central role in cell polarization processes. For example, the Rho-family GTPases, including Cdc42 (one of the most highly conserved GTPases) and its relatives (Rac in animals, Rop in plants), are crucial for controlling polarization. Polarization signals act through Cdc42 regulators to accumulate Cdc42-GTP at the future cell front. Cdc42-GTP then organizes the cytoskeleton via various effectors, shaping the polarized morphology of the cell (Chiou *et al*, 2017; Iden & Collard, 2008). Understanding the molecular details of polarity is essential for understanding various diseases, as disruptions in polarity mechanisms are often linked to cancer, neurodegenerative disorders, and immune system deficiencies (Piroli *et al*, 2019; Houston, 2017).

The discovery that bacterial cells are also polarized dates back to the first electron micrograph showing flagella at the cell poles of bacteria. Since then, efforts to study polarity-based mechanisms on model organisms including *B. subtilis, C. crescentus, and E. coli* have offered fundamental insights into the mechanisms underlying the spatiotemporal organization of bacterial cells (Kirkpatrick & Viollier, 2011; Laloux & Jacobs-Wagner, 2014; Rowlett & Margolin, 2013; Wettmann & Kruse, 2018; MacCready *et al*, 2018; Mauriello, 2019; Schumacher & Søgaard-Andersen, 2017). Similar to eukaryotes, bacterial polarity can be stably maintained over time, dictating the assembly and activity of specific cellular organelles such as flagella, pili and stalks. Alternatively, polarity can be a dynamic process involving the active diffusion and accumulation of polarized protein to precise locations as seen in the front-rear polarity of moving *Myxococcus xanthus* or *Pseudomonas aeruginosa* cells (Herrou & Mignot, 2020; Carreira *et al*, 2020; Kühn *et al*, 2023).

*M. xanthus*, a Gram-negative delta proteobacterium, glides on solid surfaces along their long axes with well-defined front-rear polarity. Directed cell motility is critical for its complex life cycle, which includes multicellular development and cooperative predation (Dinet & Mignot, 2023; Herrou & Mignot, 2020; Mercier & Mignot, 2016). *M. xanthus* cells move using two genetically distinct motility systems. The gliding (A, adventurous) motility system enables single-cell movement at the colony edges. At the molecular level, A-motility is powered by the Agl-Glt motor, which propels individual cells by adhering to specific points along the cell surface known as bacterial focal adhesions (Mignot *et al*, 2007; Faure *et al*, 2016). *M. xanthus* cells can also move in groups using a twitching-like (S, social) motility that involves type IV pili, which extend and retract to pull groups of cells together, facilitating collective movement across surfaces (Mercier *et al*, 2020). Remarkably, *M. xanthus* cells control their movement by reversing direction, achieved through the redirection of their motility apparatus to the opposite cell pole (Herrou & Mignot, 2020; Zhang *et al*, 2012a). These events termed “reversals” correspond at the cellular level, to a switching of the leading and lagging cell poles.

The establishment and maintenance of polarity reversals in *M. xanthus* involves a finely tuned interplay between signaling proteins and regulatory complexes, with the small Ras-like GTPase MglA playing a central role. Similar to other well-characterized eukaryotic small Ras-like GTPases, MglA binds to GTP (MglA_GTP_) and localizes at the leading pole where it interacts with effector proteins to activate each of the motility systems spatially, SgmX for the S-motility system (Mercier *et al*, 2020; Bautista *et al*, 2023), GltJ, AglZ and MreB for the A-motility system (Attia *et al*, 2024; Treuner-Lange *et al*, 2015). Inactivation of MglA occurs upon GTP hydrolysis because the GDP-bound state (MglA_GDP_) cannot interact with effector proteins (Dinet & Mignot, 2023; Zhang *et al*, 2010). In the cell, the MglA GTP/GDP cycle is tightly regulated by MglA interacting partners, notably the MglB protein and the complex formed by the RomR and RomX proteins (RomRX). Together, these four proteins form the core regulatory components of the polarity control network (Leonardy *et al*, 2010; Zhang *et al*, 2010, 2012b; Keilberg *et al*, 2012). The exact design of the regulatory interactions by which these proteins establish the polarity axis remains a topic of debate and is the subject of this article.

The current model suggests that the functions of MglB and RomRX are spatially separated, with MglB acting at the lagging cell pole and the RomRX complex at the leading cell pole (Herrou & Mignot, 2020; Szadkowski *et al*, 2019; Carreira *et al*, 2022). This is believed to generate a polarity axis via the following mechanism:

(i) At the lagging cell pole, MglB, a so-called roadblock domain protein is recruited by another roadblock domain protein MglC (Carreira *et al*, 2023). MglB interacts with MglA and activates GTP hydrolysis activity allosterically (see below, MglB is hence defined as a GTPase-Activating Protein (GAP) for MglA). In vitro, MglB is sufficient to activate hydrolysis, but in the cell this process is further assisted by a cognate activator, RomY (Szadkowski *et al*, 2022). *In vivo*, the action of MglB precludes accumulation of MglA at the lagging pole.

(ii) Conversely at the leading pole, RomRX is proposed to exert a dual function: first, promoting the insertion of GTP over GDP in MglA and thus acting as a guanine nucleotide exchange factor (GEF), and second, acting as a primary determinant for the polar targeting of MglA_GTP_ (Szadkowski *et al*, 2019).

This model is attractive and inspired by other polarity regulations, for example in yeast (Chiou *et al*, 2017). There are however a number of reported inconsistencies that question its validity:

- First, the mechanism of MglA polar localization is not well understood. While it is clear that an interaction between RomRX and MglA_GTP_ is required for localization at the leading cell pole, these proteins only colocalize during a short window of time after a reversal. Once the cell starts moving in the new direction, the RomRX complex relocalizes gradually at the lagging cell pole but MglA_GTP_ remains bound to the leading pole (Guzzo *et al*, 2018; Szadkowski *et al*, 2019; Herrou & Mignot, 2020). This clearly suggests that other factors maintain MglA at the leading cell pole once its localization is established. There is also the conundrum that antagonistic GAP and GEF systems co-localize at the lagging pole once polarity is established, which could only be explained by additional layers of regulations.

- Second, the GEF activity of RomRX remains questionable because conventional GEFs generally operate by first forming a complex with the GDP-bound GTPase, leading to nucleotide dissociation, and then allowing GTP-binding to displace the GEF (Bos *et al*, 2007; Cherfils & Zeghouf, 2013). In the RomRX system, RomR binds to RomX, which in turn interacts with MglA_GTP_. This ternary complex has a higher affinity for GTP, but as it cannot form in the presence of MglA_GDP_ (Szadkowski *et al*, 2019). However, GEFs generally bind to both the GTP and GDP-bound states of their substrates with equal affinity, favoring the detachment of nucleotides irrespective of their specificity, the directionality of the exchange being dependent on the relative concentrations of the nucleotides (Bos *et al*, 2007; Hennig *et al*, 2015; Cherfils & Zeghouf, 2013). Therefore, the strict interaction between RomX interacts and MglA_GTP_ is inconsistent with a GEF activity and requires a re-examination of the function of RomX and RomR *in vitro*.

- Third, although all studies converge to show that MglB activates GTP hydrolysis, its function as a canonical GAP is also questionable. First, MglB binds to MglA in a 2:1 stoichiometry, a unique configuration for GTPase/GAP complexes. This binding positions a critical arginine of MglA (Arg53) into the GTP binding site, remodeling it into a catalytic site (Baranwal *et al*, 2019; Miertzschke *et al*, 2011). While this explains how MglB promotes GTP hydrolysis, there are also a number of unexpected additional regulations that arise from the asymmetry of the MglB dimer : i) the binding of additional regulators on one of the MglB monomers, i.e. the GltJ ZnR domain (Attia *et al*, 2024), and perhaps RomY, which activates the GAP activity (Szadkowski *et al*, 2022); and ii), the association of MglA with a C-terminal helix of one of the MglB monomers (Baranwal *et al*, 2019; Chakraborty *et al*, 2024). This feature is especially intriguing because the Ct-helix prevents the dissociation of MglA_GDP_ following hydrolysis (Baranwal *et al*, 2019). This property is not canonical for GAPs, which are known to dissociate from the cognate GDP-bound GTPase (Cherfils & Zeghouf, 2013). In fact this interaction is very specific of the interaction with MglA, because MglB can also act as a GAP for SofG, another small GTPase in *M. xanthus*, but in this case it dissociates from SofG_GTP_ (Kanade *et al*, 2021). *In vivo*, the MglB Ct-helix is critical for the unipolar localization of MglB because in its absence, MglA, RomR and MglB all co-localize at both cell poles (Baranwal *et al*, 2019). The molecular mechanism for this localization pattern is not clear but it shows that regulating the interaction between MglA and MglB is essential. In summary, the Ct helix prevents the MglA-MglB complex from dissociating, which is the main basis of the current polarity model.

The observations described above call for in-depth quantitative analyses of the interactions between MglA and its regulators MglB and RomRX. In this study, we have developed a new fluorescence anisotropy assay to study these interactions. By combining these tools with an enzymatic assay that measures GTP hydrolysis, we elaborated a new enzymatic model of the MglB-dependent MglA GTP hydrolysis pathway. The model revealed the need of an additional partitioning system, which we identified as the RomRX complex. Rather than functioning as a GEF, it either interacts with MglA-GTP, acting as an effector, or prevents the formation of an MglA-MglB complex through a competitive interaction with MglB. The results suggest a model in which RomRX functions as a partitioning factor, promoting the localization of MglA at the leading cell pole immediately after reversals while preventing its localization at the lagging cell poles by interacting with MglB.

## Results

### Analysis of MglA nucleotide-binding specificities by fluorescence anisotropy

Previous work analyzed MglA nucleotide exchange *in vitro* using non-hydrolysable mant-GTP analogues (m- GMPPNP or m-GppNHp) or radioactive nucleotides (Baranwal *et al*, 2019; Chakraborty & Gayathri, 2024; Chakraborty *et al*, 2024; Zhang *et al*, 2010) of low sensitivity. The affinity of GTP for MglA could not be measured because no signal could be detected, and the affinity for GDP was not accurate because of weak and unstable signals in nucleotide exchange assays. Their binding to MglA was detected in the presence of MglA’s interacting partners; however, their binding was unstable, as the fluorescence curve showed a decrease at steady state, hindering accurate measurement of nucleotide exchange dynamics (Baranwal *et al*, 2019; Chakraborty & Gayathri, 2024; Chakraborty *et al*, 2024). We hypothesized that more sensitive fluorescent GTP/GDP derivatives, structurally compatible with MglA, could be used as alternative tools to study the dynamics of MglA nucleotide exchange both in the absence and presence of regulator proteins. Nucleotide analogues conjugated to bright dyes come in various chemistries, with fluorophores covalently bound to the base, ribose, or phosphate group of the nucleotide (Bagshaw, 2001). Because the structure of MglA bound to GTP suggests that modification of the GTP ribose should not interfere with its interaction with MglA, we selected the 2’/3’-O-(2-Aminoethyl-carbamoyl)-Guanosine-5’-tri/diphosphate GTP/GDP analogues, labeled with ATTO-488 and referred to as GTP_488_ or GDP_488_ in the remainder of the study. In these molecules, the ATTO-488 dye is attached to GTP/GDP via a long and flexible linker that allows various orientations relative to MglA and positions the dye away from the nucleotide-binding pocket **<u>(Appendix Fig.</u> <u>S1)</u>**.

The low molecular weight of these fluorescent nucleotides (MW < 1.5 kDa) relative to MglA (MW ≈ 22 kDa) and the long fluorescence lifetime of ATTO-488 (τfl = 4.1 ns) allow their binding to be monitored over time using fluorescence anisotropy (Colombo *et al*, 2021). Anisotropy measurements are not sensitive to photobleaching, which allows for the recording of signals over long periods of time. Purified MglA is found to have a GDP in its active site pocket (Baranwal *et al*, 2019). Addition of GTP_488_ or GDP_488_ to MglA (MglA_GDP_) leads to an increase of fluorescence anisotropy signals to plateau values corresponding to steady-state **(Fig. 1A)**. At initial time points, the rate limiting step of the reaction is the dissociation of GDP from MglA, which is similar in both conditions (k_off_^GDP^). The higher plateau value in the presence of GDP_488_ indicates a higher affinity of GDP_488_ compared to GTP_488_. Because MglA does not exist in a nucleotide-free state, it is only possible to determine the relative affinity of GTP_488_ or GDP_488_ for MglA. To this aim, we first incubated MglA_GDP_ with GTP_488_ or GDP_488_ and allowed the reaction to reach equilibrium and subsequently added an excess of the corresponding unlabeled nucleotide (100 µM) and monitored the decrease in fluorescence anisotropy **(Fig. 1B and C)**. The corresponding decay reaction is a measure of the dissociation rate of the fluorescent nucleotide k_off_. Fitting the data to exponential binding equations (see methods) allowed us to measure k_off_^GTP488^ = 2.8×10^-3^±5.6×10^-4^s^-1^ and k_off_^GDP488^= 6.0×10^-4^±8.7×10^-5^s^-1^, confirming the higher affinity of MglA to GDP_488_ than GTP_488_. These measurements validate our methodology; moreover, the technique is very sensitive because we could record binding at GTP_488_ concentrations as low as 10 nM **<u>(Appendix Fig. S2A)</u>.** In comparison nucleotide exchange with m-GTP could not be recorded at 100 nM (Baranwal *et al*, 2019; Chakraborty & Gayathri, 2024; Chakraborty *et al*, 2024).

**Figure 1:**
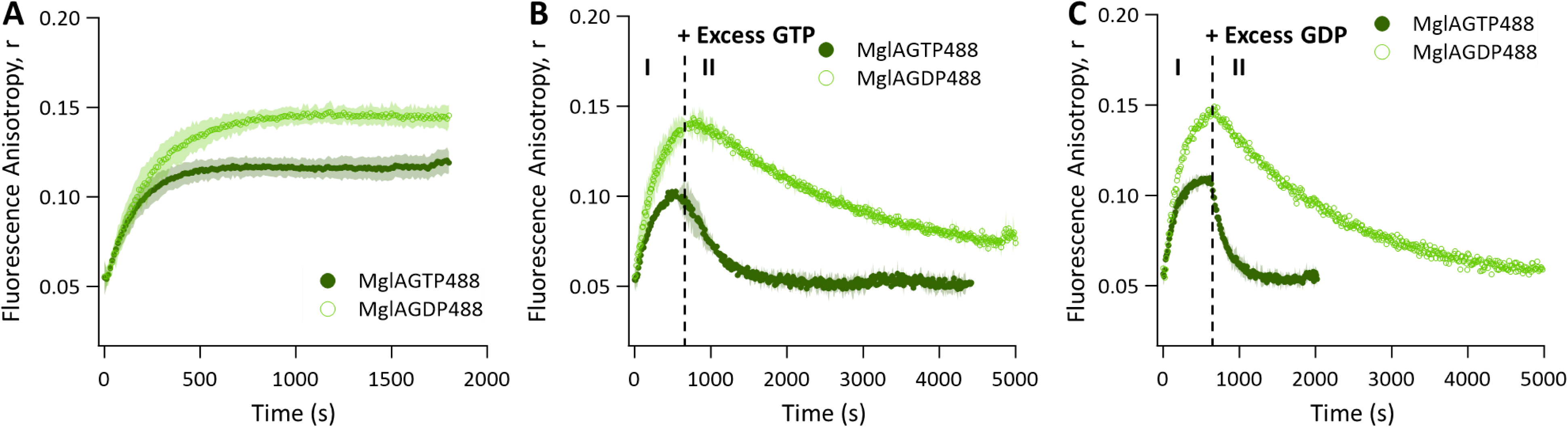
Measurement of MglA nucleotide exchange rates by fluorescence anisotropy **A.** Binding kinetics of GTP_488_ (0.1 µM; dark green) or GDP_488_ (0.1 µM; light green) in the presence of MglA_GDP_ (1 µM). **B. I-** Binding kinetics of GTP_488_ (0.1 µM; dark green) or GDP_488_ (0.1 µM; light green) in the presence of MglA_GDP_ (1 µM). **II-** After 600s, competition of GTP_488_ or GDP_488_ by addition of an excess of unlabeled GTP (100 µM) (represented by the dotted black line labeled “+ excess GTP”). **C. I-** Binding kinetics of GTP_488_ (0.1 µM; dark green) or GDP_488_ (0.1 µM; light green) in the presence of MglA_GDP_ (1 µM). **II-** After 600s, competition of GTP_488_ or GDP_488_ by addition of an excess of unlabeled GDP (100 µM) (represented by the dotted black line labeled “+ excess GDP”). The shaded regions around each curve represent the standard deviations of the fluorescence anisotropy measurement at each time point, calculated from at least three independent experimental replicates.

### RomRX does not function as a Guanine nucleotide Exchange Factor (GEF) in vitro

Next, we investigated the effect of RomRX in this experimental system. Addition of RomX led to higher values of anisotropy at steady-state in the presence of GTP_488_, but not in the presence of GDP_488_. This result indicates an increase in the molecular weight of MglA_GTP488_ specifically, due to its binding to RomX. In contrast, RomR did not interact directly with either form of MglA **(Fig. 2A-B)**. When RomR and RomX were added together with MglA_GTP488_, a ternary complex formed as evidenced by an additional increase in the fluorescence anisotropy signal **(Fig. 2A)**. The binding mode of the RomRX complex is thus clearly different from that expected for a GEF and more similar to the binding of an MglA effector. To test this directly, we compared RomX binding to that of a known MglA effector, the Ct Tetratricopeptide Repeat triad of the SgmX Type-IV pilus activator (SgmX-Ct) (Mercier *et al*, 2020). Similarly, SgmX-Ct binds only to MglA_GTP488_ but not to MglA_GDP488_ **(Fig. 2C-D)**. We could determine that MglA_GTP488_ binds RomX with a K_D_ = 1.7 ± 0.3 µM, which was slightly decreased to K_D_ = 1.0 ± 0.4 µM upon addition of RomR **(Fig. 2E-F)**. MglA_GTP488_ binds SgmX-Ct with a slightly lower K_D_ = 0.6 ± 0.3 µM, but in the same order of magnitude **(Fig. 2G)**. Thus, MglA_GTP488_ binds RomX and SgmX-Ct by similar modes.

**Figure 2:**
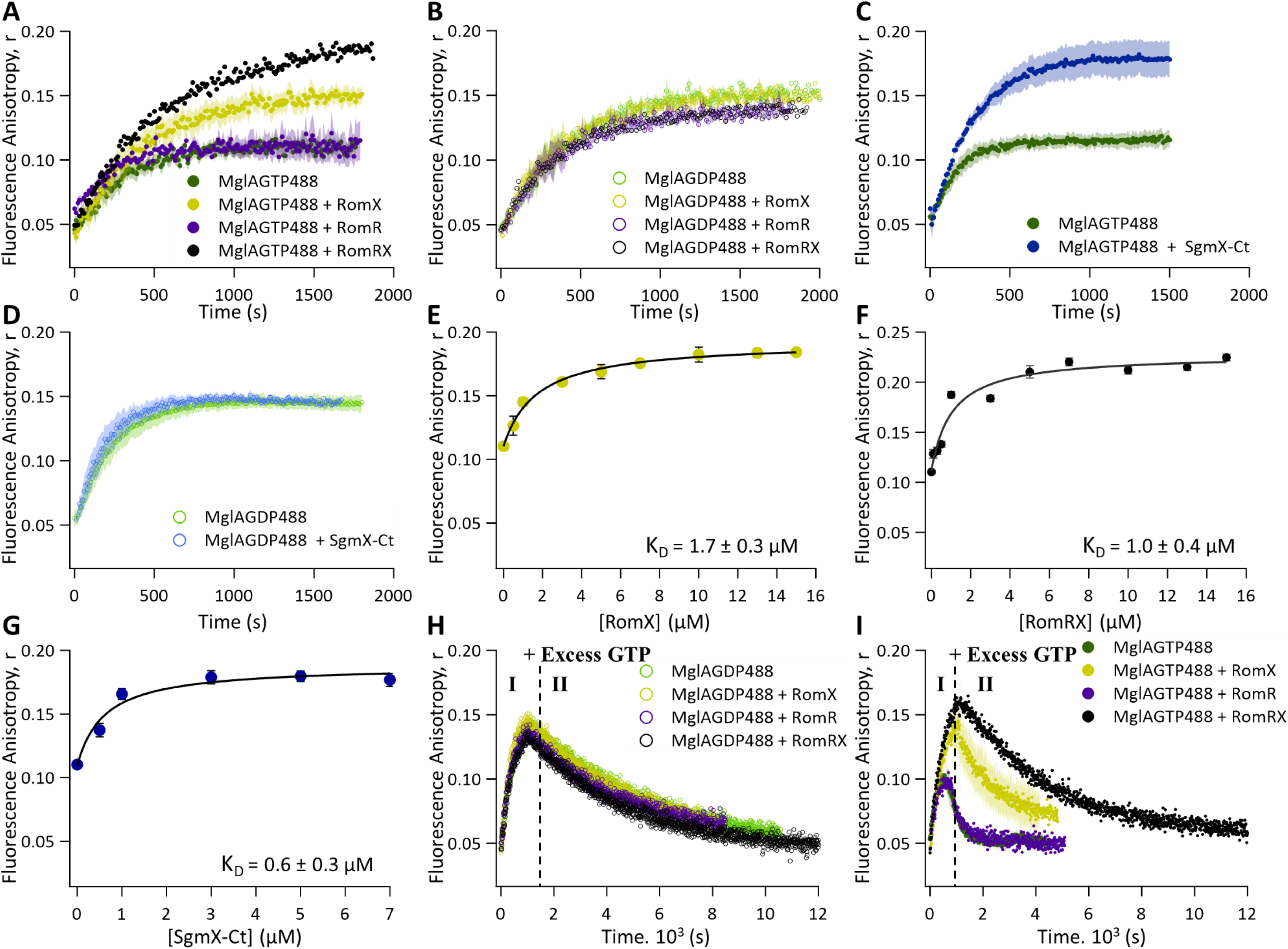
RomRX complex acts as an MglA effector. **A.** Binding kinetics of GTP_488_ (0.1 µM) to MglA_GDP_ (1 µM) in the absence (green) or in the presence of RomX (1µM; yellow), RomR (1µM; purple) or RomRX (1µM; black). **B.** Binding kinetics of GDP_488_ (0.1 µM) to MglA_GDP_ (1 µM) in the absence (green) or in the presence of RomX (1µM; yellow), RomR (1µM; purple) or RomRX (1µM; black). **C.** Binding kinetics of GTP_488_ (0.1 µM) to MglA_GDP_ (1 µM) in the absence (green) or in the presence of SgmX-Ct (1µM; blue). **D.** Binding kinetics of GDP_488_ (0.1 µM) to MglA_GDP_ (1 µM) in the absence (green) or in the presence of SgmX-Ct (1µM; blue). **E-G.** Fluorescence anisotropy measurements for RomX, RomRX or SgmX-Ct respectively binding to MglA_GTP488_. Binding was estimated by measuring the change in fluorescence anisotropy of MglA_GTP488_ upon titration with increasing concentrations of RomX, RomRX or SgmX-Ct. Each reading corresponds to the value of fluorescence anisotropy (30 min after the start of the experiment, at steady state). Each point represents the mean value from at least three independent measurements, and the error bars indicate standard deviation. **H. I-** Binding kinetics of GDP_488_ (0.1 µM) to MglA_GDP_ (1 µM) in the absence (green) or in the presence of RomX (1µM; yellow), RomR (1µM; purple) or RomRX (1 µM, black). **II-** After 600s, competition of GDP_488_ by addition of an excess of unlabeled GTP (100 µM) (represented by the dotted black line labeled “+ excess GTP”). **I. I-** Binding kinetics of GTP_488_ (0.1 µM) to MglA_GDP_ (1 µM) in the absence (green) or in the presence of RomX (1µM; yellow), RomR (1µM; purple) or RomRX (1 µM, black). **II-** After 600s, competition of GTP488 by addition of an excess of unlabeled GTP (100 µM) (represented by the dotted black line labeled “+ excess GTP”). The shaded regions around each curve in panels A-D, H and I represent the standard deviations of the fluorescence anisotropy measurements at each time point, calculated from at least three independent experimental replicates.

Small G-protein effectors tend to stabilize the GTP-bound state of the cognate G-protein, which might explain why RomRX was proposed to facilitate nucleotide exchange (Szadkowski *et al*, 2019). If this was correct, this effect should also be observed with SgmX-Ct. We tested that possibility by conducting chase experiments, preloading MglA with GDP_488_ or GTP_488_ and then adding an excess of unlabelled GTP to monitor rates of nucleotide release. We did not detect any effect of RomX, RomR or RomRX on the rate of GDP_488_ to GTP exchange **(Fig. 2H)** but we observed that binding to RomX stabilizes the binding of GTP_488_ on MglA **(Fig. 2I)**. In the presence of RomX, we measured a k_off_^GTP488^ = 8.4×10^-4^ ± 3.3×10^-4^s^-1^, compared to 2.8×10^-3^±5.6×10^-4^s^-1^ in its absence **(Fig. 2I and Table 1)**. This effect is even more pronounced in the presence of RomRX (k_off_^GTP488^ = 2.4×10^-4^ ± 5.5×10^-5^ s^-1^). Most importantly, SgmX-Ct also stabilized MglA_GTP488_ and to similar levels (k_off_^GTP488^ = 2.7×10^-4^ ± 2.4 ×10^-5^s^-1^) **(<u>Appendix Fig. S2B</u> and Table 1)**. Thus, we conclude that RomX and the RomRX complex are *bona fide* MglA effectors like SgmX-Ct and not an MglA GEF as originally proposed.

**Table 1:** Dissociation rates constant (K_off_) corresponding to the graph in Fig .2I. The errors represent the standard deviations calculated from at least three independent measurements.

|  |  |
| --- | --- |
| | $k_{\text{GTP488}}^{\text{off}} \text{ (s}^{-1}\text{)}$ |
| **MglA<sub>GDP</sub>** | $2.8 \times 10^{-3} \pm 5.6 \times 10^{-4}$ |
| **MglA<sub>GDP</sub> + SgmX-Ct** | $2.7 \times 10^{-4} \pm 2.4 \times 10^{-5}$ |
| **MglA<sub>GDP</sub> + RomX** | $8.4 \times 10^{-4} \pm 3.3 \times 10^{-4}$ |
| **MglA<sub>GDP</sub> + RomR** | n.r. |
| **MglA<sub>GDP</sub> + RomRX** | $2.4 \times 10^{-4} \pm 5.5 \times 10^{-5}$ |

### MglB is an atypical MglA GTPase-activating protein

We next used the anisotropy assay to explore the interaction between MglA and MglB. Addition of GTP_488_ to a mix of MglA_GDP_ and MglB led to higher anisotropy levels than observed in the absence of MglB, demonstrating that an MglA_GTP488_-MglB complex is formed **(Fig. 3A)**. However, after reaching a maximal value, the anisotropy signal decreased with slow kinetics. This decrease must correspond to the hydrolysis of the GTP_488_ nucleotide bound to MglA into GDP_488_ driven by the GAP activity of MglB, because it was not observed when an MglA mutant (MglA^Q82L^) that is resistant to hydrolysis was co-incubated with MglB **(<u>Appendix Fig. S2C and D</u>)**. The persistently high anisotropy signal also suggests that MglA_GDP488_ remains attached to MglB. To prove this, we incubated MglA_GDP_ with MglB and added GDP_488_. We observed a sharp increase in the anisotropy signal indicating that MglB also interacts with MglA_GDP488_ **(Fig. 3A)**. The signals observed with MglA_GTP488_ and MglA_GDP488_ converged over time, confirming an MglB-induced GTP hydrolysis **(Fig. 3A inset)**. Thus, consistent with findings by Baranwal et al., MglB behaves as a non-typical GAP, activating GTP hydrolysis but not dissociating from MglA_GDP_. This result was further confirmed with an MglB version lacking the Ct helix (MglB^ΔCt^), which bound MglA_GTP488_ but not MglA_GDP488_ **(<u>Appendix Fig.</u> <u>S2E and F</u>)**.

**Figure 3:**
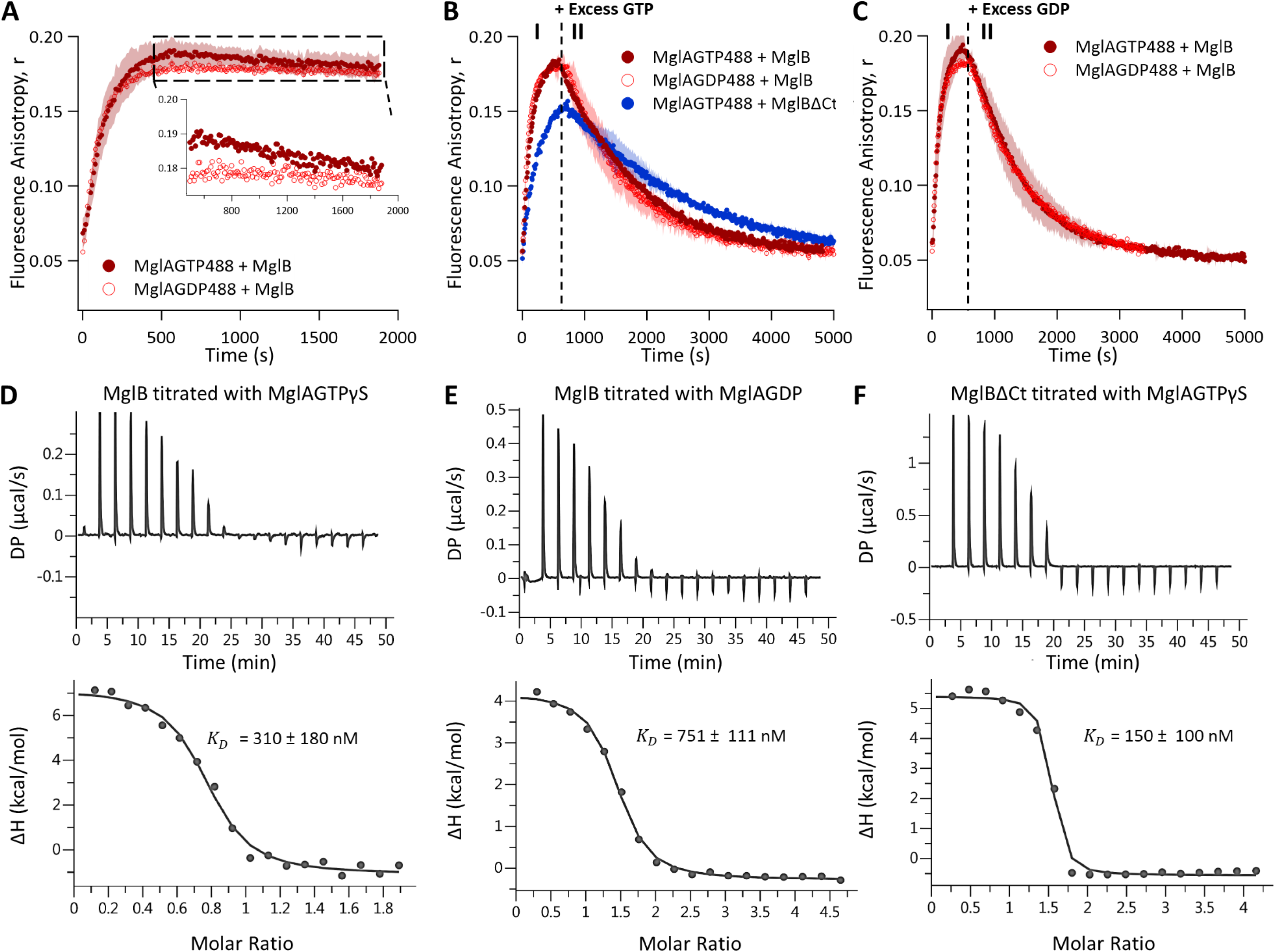
Characterization of MglA interactions with MglB and its mutants **A.** Binding kinetics of GTP_488_ (0.1 µM; dark red) or GDP_488_ (0.1 µM; light red) to MglA_GDP_ (1 µM) in the presence of MglB (2 µM). The inset corresponds to a zoomed view of the dashed rectangle in the graph. **B. I-** Binding kinetics of GTP_488_ (0.1 µM; dark red) or GDP_488_ (0.1 µM; light red) to MglA_GDP_ (1 µM) in the presence of MglB (2 µM) or MglB^ΔCt^ (blue; 2 µM). **II-** Competition of GTP_488_ or GDP_488_ by addition of an excess of unlabeled GTP (100 µM) at 600s (represented by the dotted black line labeled “+ excess GTP”). **C. I-** Binding kinetics of GTP_488_ (0.1 µM; dark red) or GDP_488_ (0.1µM; light red) to MglA_GDP_ (1 µM) in the presence of MglB (2 µM). **II-** Competition of GTP_488_ or GDP_488_ by addition of an excess of unlabeled GDP (100 µM) at 600s (represented by the dotted black line labeled “+ excess GDP”). The shaded regions around each curve in panels A-C represent the standard deviations of the fluorescence anisotropy measurements at each time point, calculated from at least three independent experimental replicates. Raw (Top panel) and integrated heat (bottom panel) plots for the titration of: **D.** 200 µM MglB into 20 µM MglA and 100 µM GTPγS **E.** 492 µM MglB into 20 µM MglA and 100 µM GDP **F.** 1.1mM MglB^ΔCt^ into 50 µM MglA and 100 µM GTPγS. The binding isotherms are a representation of replicate experiments.

MglB has also been proposed to facilitate GTP exchange upon binding to MglA (Baranwal *et al*, 2019; Chakraborty *et al*, 2024), which might again be due to an effector-type interaction. To verify this result, we performed chase experiments with unlabeled nucleotides in the presence of MglB or MglB^ΔCt^ **(Fig. 3B and C)**. Consistent with previous observations, we found that MglA exhibits enhanced affinity for GTP_488_ than for GDP_488_ in the presence of MglB (k_off_^GTP488^ = 9.0×10^-4^ ± 1.0×10^-4^ s^-1^ and k_off_^GDP488^ = 1.2×10^-3^ ± 2.6×10^-4^ s^-1^) **(Table 2)**. We also used isothermal titration calorimetry (ITC) to measure MglA affinities for MglA_GTPγS_ (a non-hydrolysable form of MglA_GTP_) and MglA_GDP_ in the presence of MglB. We observed as expected a lower dissociation constant K_D_ = 310 ± 180 nM for MglA_GTPγS_ than for MglA_GDP_, where K_D_ = 751 ± 111 nM **(Fig. 3D and E)**. Furthermore, the Ct helix of MglB stabilizes the MglA_GTP_-MglB complex, favoring GTP binding **(Fig. 3B and Table 2).** This stabilization is further evidenced by the lower K_D_ of 150 ± 100 nM for the interaction between MglB^ΔCt^ and MglA_GTPγS_ **(Fig. 3F)**.

**Table 2:**
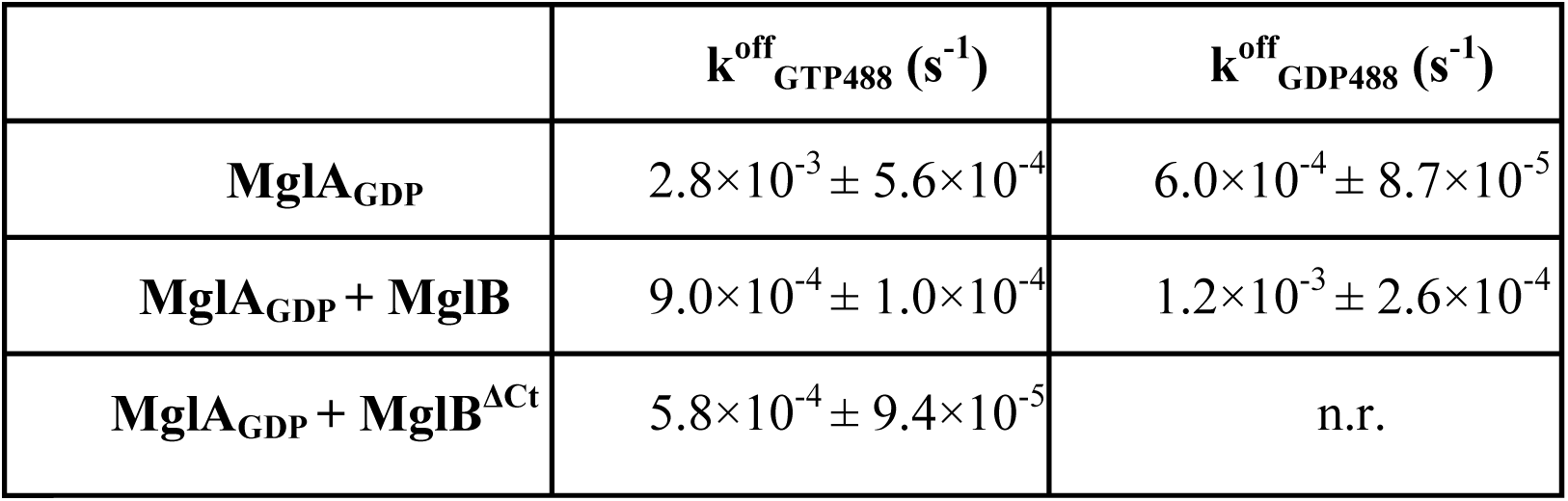
The dissociation rates constant (k_off_) corresponding to the graph in. **Fig. 3B and C. The errors represent the standard deviations calculated from at least three independent measurements.**

### A new enzymatic model of MglA and MglB interaction

The results presented above highlight the function of RomRX as an MglA effector and confirm that MglB does not dissociate from MglA_GDP_, which deeply questions the prevailing model. Similarly, MglB cannot be called a GEF (Baranwal *et al*, 2019; Chakraborty *et al*, 2024) because it does not dissociate from MglA_GTP_ to which it binds with high affinity. Thus, we developed an enzymatic model based on the current knowledge of the system to elucidate the exact properties of the MglA-MglB complex. In this model, MglA binds with GTP or GDP, forming a binary complex. Each of these complexes can also bind to MglB, forming MglAGTP- MglB or MglA_GDP_-MglB ternary complexes. Each ternary complex can spontaneously exchange nucleotides, but also hydrolyse GTP leading to the release of inorganic phosphate (Pi) without complex dissociation **(Fig. 4A).**

**Figure 4:**
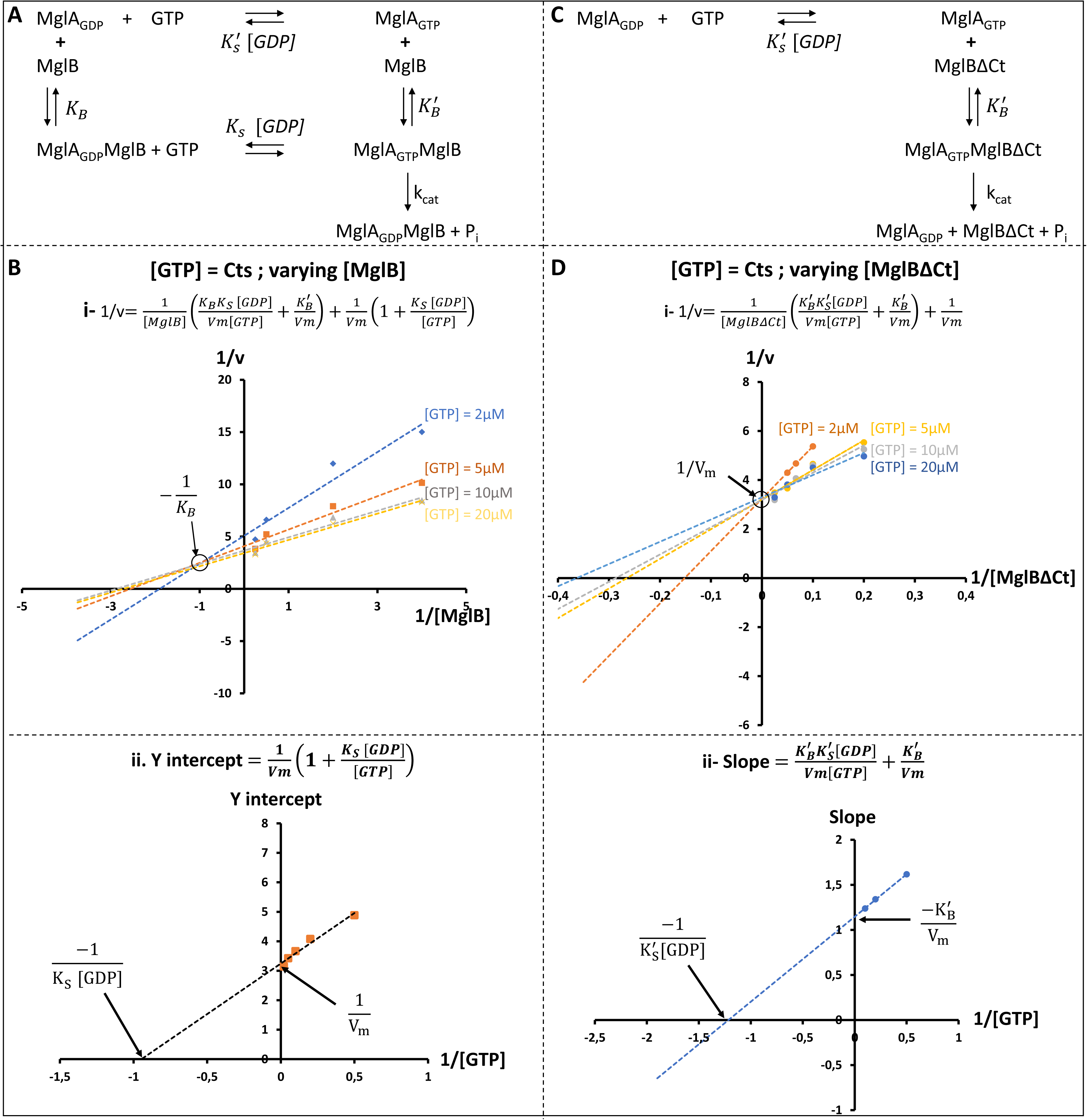
Kinetic analysis of MglA’s interaction with MglB or MglB^ΔCt^ in the presence of GTP. **A.** Schematic representation of the rapid equilibrium random fixation model describing MglA and MglB kinetic interaction. **B. Top:** Plot of 1/v as a function of 1/[MglB] at varying GTP concentrations (2, 5, 10 and 20µM). The data were modeled using the equation (i). All lines converge to a single point corresponding to -1/K_B_. **Bottom:** Following equation (ii), the y-intercept (corresponding to the plot of 1/v in the function of 1/[MglB] at a single GTP concentration) was plotted in the function of 1/[GTP] at four different GTP concentrations 5, 10,20 and 50 µM. The Y-intercept corresponds to 1/Vm and the x-intercept corresponds to -1/K_s_[GDP]. **C.** Schematic representation of the rapid equilibrium ordered fixation model describing MglA and MglB^ΔCt^ kinetic interaction **D. Top:** Plot of 1/v as a function of 1/[MglBΔCt] at varying GTP concentrations (2, 5, 10 and 20µM). The data were modeled using the equation (i). All lines converge to a single point corresponding to 1/V_m_. **Bottom:** Following equation (ii), the slope (corresponding to the plot of 1/v in the function of 1/[MglBΔCt] at a single GTP concentration) was plotted in the function of 1/[GTP] at three different GTP concentrations 2, 5 and 10 µM. The Y-intercept corresponds to -K ^’^/Vm and the x-intercept corresponds to -1/K ^’^[GDP].

The differential equations describing the model are consistent with a rapid equilibrium random fixation mechanism (Ivanetich & Goold, 1989) **(<u>Extra Methods</u>).** The term “rapid equilibrium” indicates that the substrates and products binding and release occur much faster than the catalytic conversion step. The system involves two reactants (i.e. the substrate; GTP and the activator; MglB) and two products (i.e. P_i_ and GDP). However, in our case GDP remains in the nucleotide pocket until the second round of fixation. This model predicts that varying [GTP] or [MglB] results in linear regressions in double reciprocal Lineweaver-Burk (LW) plots (1/ν vs. 1/[MglB] or 1/[GTP]) consistent with Henri-Michaelis-Menten kinetics, intersecting at the same value left of the 1/ν axis. Secondary plots of the y intercepts and slopes from the primary lineweaver-burk plots, plotted as function of 1/[GTP] or 1/[MglB], should also be linear. To test this model experimentally, we measured GTP hydrolysis by determining Pi release kinetics using an established enzyme-coupled assay (Webb, 1992) and constructed LW plots and their secondary plots by varying [MglB] or [GTP] **(Fig. 4B and <u>EV1A</u>)**. The obtained experimental curves aligned remarkably well with the rapid equilibrium random fixation model for the MglA-MglB complex, allowing determination of the system constants **(Fig. 4B**, **Table 3)**.

**Table 3:**
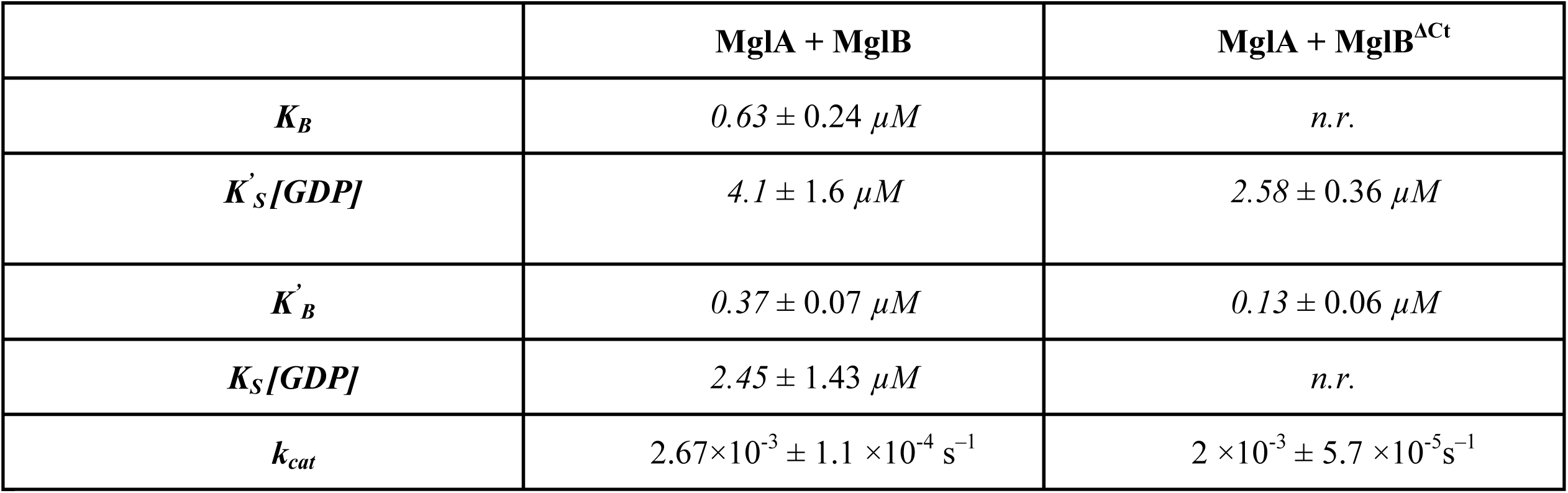
Rapid equilibrium random/ordered mechanism. Kinetic parameters obtained from solving the graphs in. **Fig. 4B and D and <u>Fig. EV1A and B</u>**

To further understand the function of the MglB Ct helix, we also constructed a model analyzing the behavior of the system when this helix is absent, making the interaction of MglB with MglA specific to its GTP-bound form **(<u>Appendix Fig. S2E and F</u>)**. In this scenario, the system becomes coherent with a rapid equilibrium ordered fixation mechanism (Posner *et al*, 1992), where the MglA-MglB^ΔCt^ ternary complex forms only upon interaction with MglA_GTP_. This leads to the dissociation of MglA_GDP_ from MglB following hydrolysis **(Fig. 4C)**. The differential equations for this model are shown in **<u>Extra Methods</u>** and the LW plots in **Fig. 4D** and **EV1B**. The LW plots differ from the random model when 1/v is plotted against 1/[MglB^ΔCt^] at various fixed GTP concentrations, intersecting on the vertical axis **(Fig. 4D)**. This indicates no velocity dependence on MglB^ΔCt^ concentration at saturating GTP, similar to previous observation (Baranwal *et al*, 2019) and characteristic of a rapid equilibrium ordered mechanism with GTP binding before MglB^ΔCt^.

The kinetic parameters from the two enzymatic models are remarkably consistent with the affinity values measured by ITC as one falls within the uncertainty range of the other. (compare affinities calculated for MglA_GTP_ and MglB K^’^ = 370 ± 70 nM *vs* 310 ± 180 nM by ITC, and for MglA_GDP_ and MglB, K_B_ = 630 ± 240 nM vs 751± 111 nM by ITC). In addition, the model is also consistent with a stabilizing effect of MglB on MglA-GTP (compare K_S_ [GDP] = 2.45 ± 1.43 µM in the presence of MglB to K^’^ [GDP] = 4.1 ± 1.6 µM in its absence), following the same trend as in the anisotropy experiments. The measurements also confirm that the deletion of the MglB Ct helix increases the affinity to MglA-GTP (compare K^’^ = 370 ± 70 nM for MglB^ΔCt^ versus K^’^ = 130 ± 100 nM for MglB, which is very similar to K_D_ = 150 ± 100 nM, measured by ITC). The measured k_cat_ values were similar for MglB^ΔCt^ (2 ×10^-3^ ± 5.7 ×10^-5^s^−1^) compared to MglB (2.67×10^-3^ ± 1.1 ×10^-4^ s^−1^), which are both very low compared to canonical GAPs. For example, p120 GAP stimulates the GTPase reaction of Ras with a k_cat_ ∼19 s^-1^ under saturating conditions (Wittinghofer *et al*, 1997). This is consistent with measurements by Galicia et al. (Galicia *et al*, 2019) and suggests that MglB may require interacting proteins to fully activate or enhance its GAP activity (ie RomY, see discussion).

Taken together, our results demonstrate that MglA-RomRX and MglA-MglB interactions do not correspond to canonical GEF and GAP interactions. The enzymology model above should therefore be taken into consideration in any new hypothesis attempting to explain their function in cell polarization (See below and Discussion)

### RomR acts on MglB to inhibit the formation of the MglA-MglB complex

*In vivo*, MglB is required to exclude MglA from the lagging pole, which was previously assumed to be linked to its GTPase activity. The observation that MglA, whether bound to GTP or GDP, forms a strong complex with MglB that does not dissociate upon GTP hydrolysis raises the question of the involvement of additional regulators. We first tested how effectors such as SgmX and RomX compete with formation of the MglA-MglB complex. We found that even at high concentrations (up to 30 µM), these proteins were not able to dissociate the complex **(Fig. 5A and <u>Appendix Fig. S2G-J</u>).**

**Figure 5:**
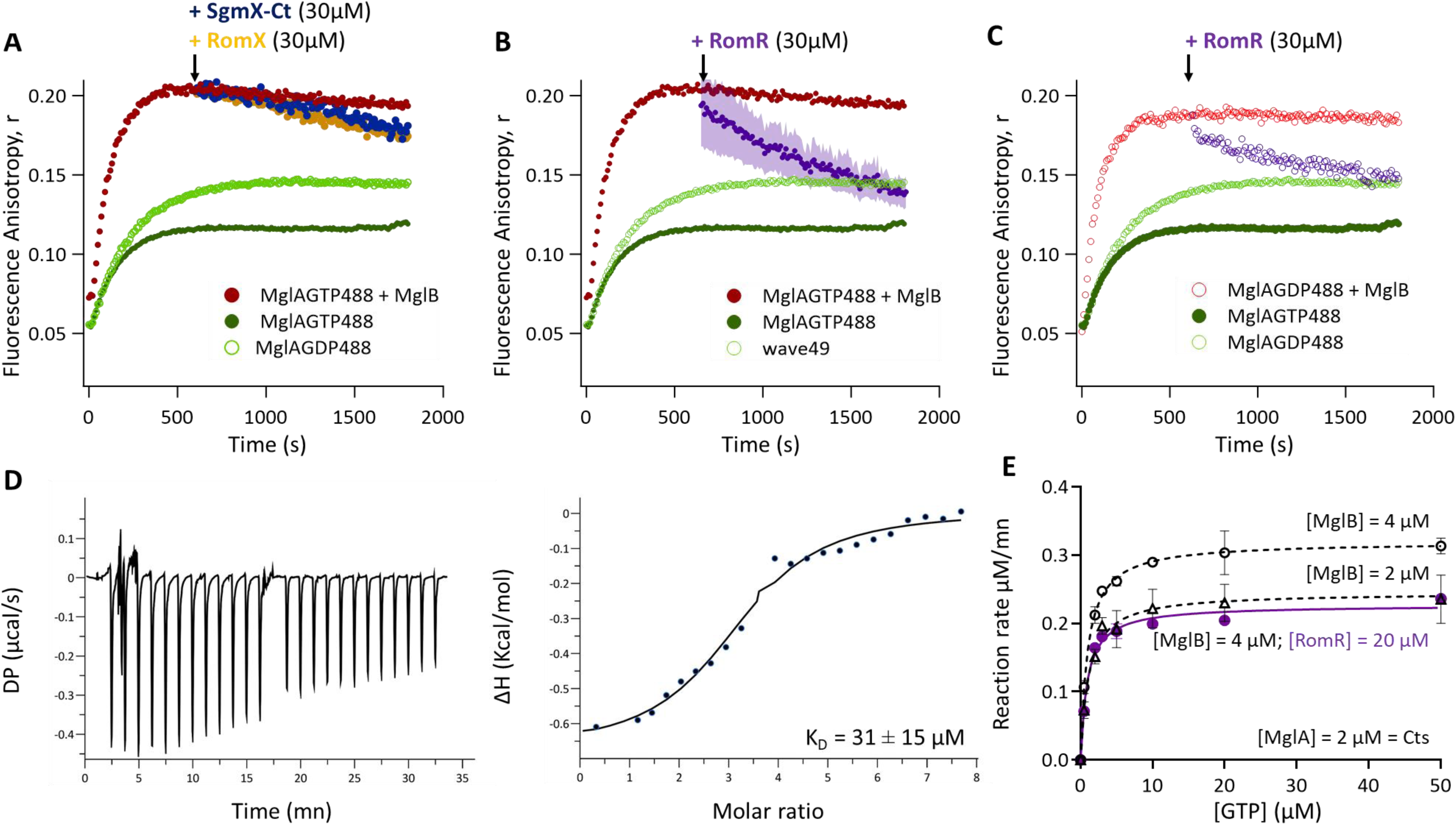
Impact of the RomR on the MglA-MglB interaction **A.** Binding kinetics of GTP488 (dark green;0.1 µM) or GDP488 (light green; 0.1 µM) to MglA_GDP_ (1 µM). Binding kinetics of GTP488 (0.1 µM) to MglA_GDP_ in the presence of MglB (2µM; dark red) or binding kinetics of GTP488 (0.1 µM) to MglA_GDP_ (1 µM) in presence of MglB (2µM; dark red) followed by the addition of RomX (30 µM; orange) or SgmX-Ct (30 µM; dark blue) at 600s. **B.** Binding kinetics of GTP488 (dark green;0.1 µM) or GDP488 (light green; 0.1 µM) to MglA_GDP_ (1 µM). Binding kinetics of GTP488 (0.1 µM) to MglA_GDP_ in the presence of MglB (2µM; dark red) or binding kinetics of GTP488 (0.1 µM) to MglA_GDP_ (1 µM) in presence of MglB (2µM; dark red) followed by the addition of RomR (30 µM; purple) at 600s. **C.** Binding kinetics of GTP488 (dark green;0.1 µM) or GDP488 (light green; 0.1 µM) to MglA_GDP_ (1 µM). Binding kinetics of GDP488 (0.1 µM) to MglA_GDP_ in the presence of MglB (2µM; light red) or binding kinetics of GDP_488_ (0.1 µM) to MglA_GDP_ (1 µM) in presence of MglB (2µM; light red) followed by the addition of RomR (30 µM; purple) at 600s. **D.** Isothermal titration calorimetry (ITC) profile of MglB-RomR interaction. Raw (left panel) and integrated heat (right panel) plots for the titration of 1.3 mM MglB into 70 µM RomR. The fit yielded a K_D_ of 31 ± 15 µM. **E.** Initial rate data for Pi release at different concentration of GTP in presence of MglA_GDP_ (2 µM) + MglB (4 µM) (empty circle), MglA_GDP_ (2 µM) + MglB (2 µM) (empty triangle) or MglA_GDP_ (2 µM) + MglB (4 µM) + RomR (20 µM) (purple filled square).

Thus, effector competition may not be sufficient to dissociate MglA-MglB. Given that RomR localizes at the lagging pole and directly interacts with MglB (although there are conflicting evidence for this interaction, which has recently been challenged by the same group (Keilberg *et al*, 2012; Carreira *et al*, 2023)), we tested whether the addition of RomR affects the MglA_GTP_-MglB complex. Addition of 30 µM RomR led to an abrupt decrease in fluorescence anisotropy signal **(Fig. 5B)** indicating that RomR triggers the dissociation of the complex. Adding 30µM RomR also disrupted the MglA_GDP_-MglB complex **(Fig. 5C)**. Since RomR does not interact with MglA, it likely competes with MglA for binding to MglB, thereby dissociating the complex and reducing the available MglB for MglA interaction.

There is conflicting data regarding a potential direct interaction between RomR and MglB, as some pull-down experiments detect this interaction while others do not (Keilberg *et al*, 2012; Carreira *et al*, 2023). We revisited this potential direct interaction using ITC and found a low affinity interaction with a K_D_ of 31 ± 15 µM **(Fig. 5D)**. Microscale thermophoresis also confirmed these interactions, giving a K_D_ value in the same order of magnitude (∼ 59 µM, <u>Appendix Fig. S3A</u>).

Previous studies (Szadkowski *et al*, 2019) have shown that RomR does not affect the GTPase activity of MglA when added at equal concentration to MglB. However, due to the low affinity interaction between RomR and MglB, significant changes in activity should not be expected unless RomR is present at high concentrations. To test this prediction, we conducted GTPase assays in the presence of varying concentrations of MglB and a fixed concentration of RomR (20 µM) **(Fig. 5E and** <u>Appendix Fig. S3B and C</u>). Michaelis-Menten and Lineweaver-Burk plots show that RomR significantly reduced the reaction rate, mirroring the effect observed at lower effective concentrations of MglB (an approximately twofold decrease in enzyme efficiency *k*^*app*^_*cat*_/*k*^*app*^_*m*_) **(Fig. 5E and Table 4)**. This shows that when the concentration of RomR is close to the dissociation constant of the MglB-RomR complex, it effectively competes with the formation of the MglA-MglB complex.

**Table 4:** Kinetic parameters table. Initial rate data were fitted to the Michaelis-Menten equation. The assays were performed using constant concentration of proteins and a range of GTP concentration (0.5 to 100 µM).

| | $k_{\text{cat}}^{\text{app}} (\text{min}^{-1})$ | $K_m^{\text{app}} (\mu\text{M})$ | $k_{\text{cat}}^{\text{app}}/K_m^{\text{app}} (\text{M}^{-1}\text{s}^{-1})$ |
| --- | --- | --- | --- |
| **MglA (2$\mu\text{M}$) + MglB (4$\mu\text{M}$)** | $0.16 \pm 0.011$ | $1.02 \pm 0.014$ | $2.60 \times 10^3$ |
| **MglA (2$\mu\text{M}$) + MglB (2$\mu\text{M}$)** | $0.12 \pm 0.001$ | $0.98 \pm 0.22$ | $2.04 \times 10^3$ |
| **MglA (2$\mu\text{M}$) + MglB (4$\mu\text{M}$) - RomR (20$\mu\text{M}$)** | $0.11 \pm 0.002$ | $1.00 \pm 0.023$ | $1.83 \times 10^3$ |

### In vitro reconstitution of MglA exclusion by the RomR-MglB complex in a cell-like configuration

The results above suggest that RomR modulates the interaction between MglA and MglB and thus controls the turnover rate of the MglA-MglB complex *in vivo*. However, because the affinity between MglB and RomR is low, this effect is only expected if there is sufficient accumulation of RomR to compete with the formation of the high affinity MglA-MglB complex. Such accumulation is possible in vivo because most of the RomR pool relocalizes with RomX to the lagging pole (**Fig. 6A and B**). To test this possibility in conditions that mimic the cellular context closely, we formed individual spherical droplets of 10 to 20 µm- diameter that reproduce the concave shape of the bacterial poles, although the radii of curvature are larger and constant around the droplets **(Fig. 7A and <u>Appendix Fig. S4A</u>)**.The lipids stabilizing the droplet interface were selected to be similar in composition to the inner membrane of *M. xanthus*. We fluorescently labeled MglA, MglB, and RomR. MglA was either labeled directly (**<u>Appendix Fig. S4B</u>**) or its localization was tracked using bound fluorescent nucleotides as a label **(Fig. 7B-C and <u>Appendix Fig. S4C</u>)**. As shown by the anisotropy measurements, fluorescent nucleotides remain stably bound to MglA for at least 20 minutes, justifying colocalization of fluorescent nucleotides and MglA for the duration of these experiments. MglA (in its GTP or GDP states) and RomR localized diffusely in these compartments at any concentration **(Fig. 7B-D and <u>Appendix Fig. S4B-C and S5A</u>)**. In contrast, MglB localized in large domains formed at the lipid/water interface **(Fig. 7E and <u>Appendix Fig. S5B</u>)**. Control experiments in the absence of lipids did not reveal any MglB binding (**<u>Appendix Fig. S6</u>**). This observation is consistent with previous work showing that MglB binds weakly to liposomes (Galicia *et al*, 2019). The characteristic localization of MglB in the droplet provided a good opportunity to characterize protein-protein interactions in this cell-like configuration using two-color imaging.

**Figure 6:**
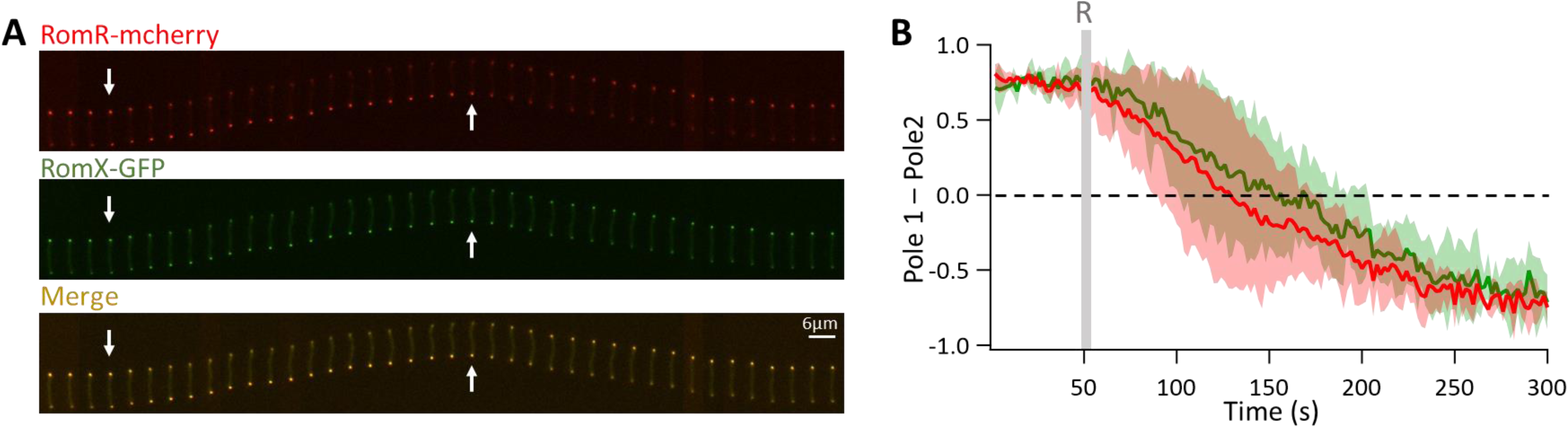
In vivo localization and dynamics of RomR and RomX. A. Time-lapse fluorescence microscopy images of pole-to-pole dynamics of RomR-mcherry (red) and RomX- GFP (green) (20s intervale). The merged image highlights colocalization of both proteins at polar regions. Arrows point to reversal events. The scale bar represents 6 μm. B. Quantitative analysis of fluorescence intensity dynamics over time, comparing the localization of RomR-mCherry and RomX-GFP from Pole 1 to Pole 2. The difference in intensities of RomR-mCherry and RomX- GFP between the two poles was normalized to the range defined by the minimum and maximum intensity. Shaded regions represent the standard deviation across measurements from 3 independent cells. The 0 on the y-axis indicates symmetrical localization between the two poles. The gray line indicates the time of reversal.

**Figure 7:**
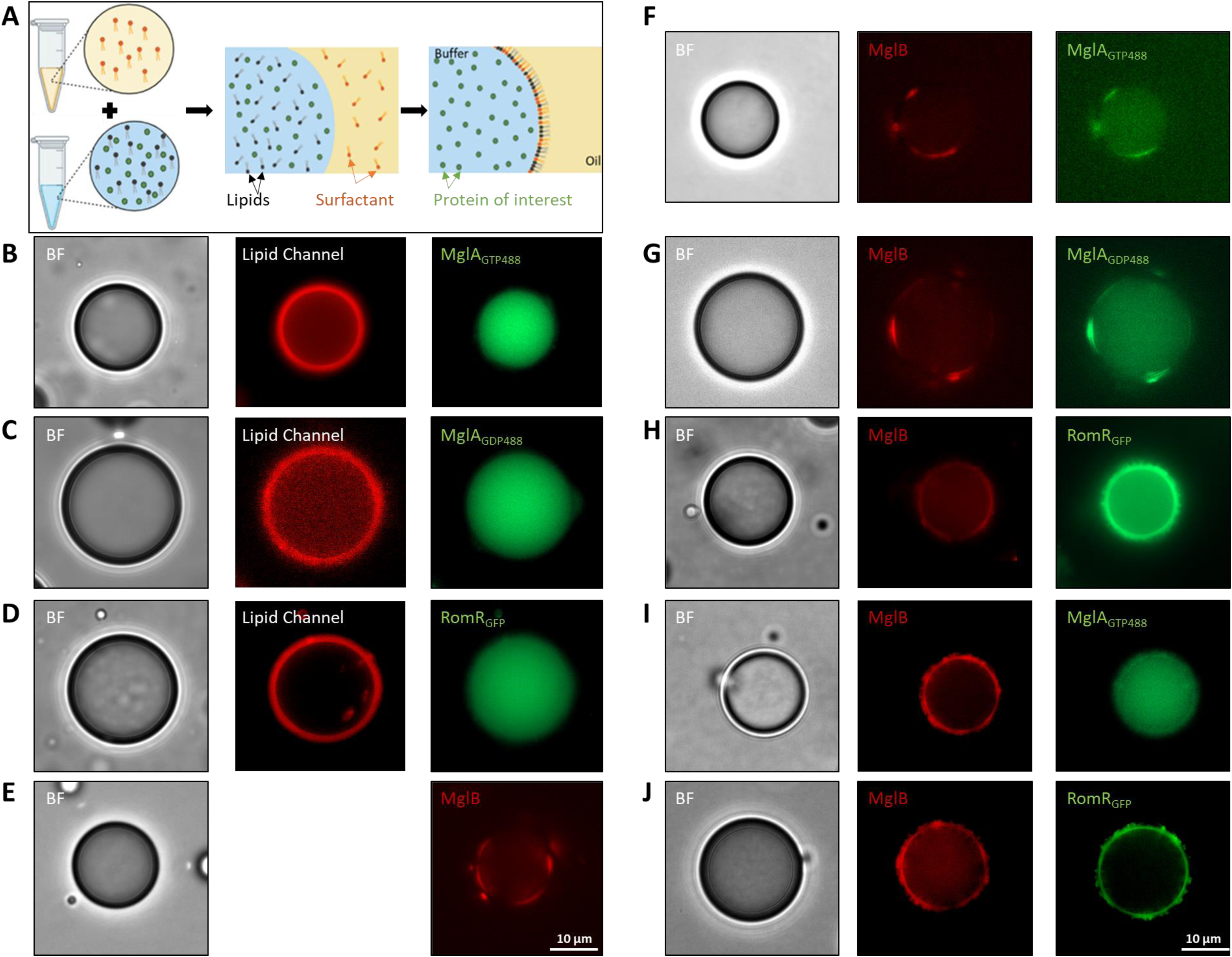
Epi-fluorescence imaging of MglA, MglB, and RomR localization in water-in-oil emulsion droplets stabilized by lipids **A.** Schematic representation of the formation of the lipid droplets; surfactants are in orange, lipids are in black and proteins are in green. **B.** Epi-fluorescence image of lipid droplets containing MglA (1 µM) labeled with GTP_488_ (0.2 µM) **C.** Epi-fluorescence image of lipid droplets containing MglA (1 µM) labeled with GDP_488_ (0.2 µM) **D.** Epi-fluorescence image of lipid droplets containing RomR-GFP (10 µM). **E.** Epi-fluorescence image of lipid droplets containing red-labeled MglB (2 µM). **F.** Epi-fluorescence images of lipid droplets containing MglA (1 µM) labeled with GTP_488_ (0.2 µM) in presence of red-labeled MglB (2 µM). **G.** Epi-fluorescence images of lipid droplets containing MglA (1 µM) labeled with GDP_488_ (0.2 µM) in presence of red-labeled MglB (2 µM). **H.** Epi-fluorescence images of lipid droplets containing red-labeled MglB (10 µM) in presence of RomR-GFP (25 µM). **I.** Epi-fluorescence images of lipid droplets containing red-labeled MglB (10 µM) in presence of RomR-GFP (25 µM) and unlabeled MglA (5 µM) **J.** Epi-fluorescence images of lipid droplets containing red-labeled MglB (10 µM) in presence of unlabeled RomR (25 µM) and MglA (5 µM) labeled with GTP_488_ (0.2 µM). Rhodamine was used to label the lipids.

Mixing red-labeled MglB with green-labeled MglA_GTP488_ or MglA_GDP488_ did not modify the localization of MglB but led to the recruitment of MglA at the lipid/water interface with MglB **(Fig. 7F-G)**. This observation demonstrates that MglB and MglA interact in the droplet assay and that this interaction does not interfere with MglB lipid-binding. Mixing red-labeled MglB with RomR-GFP also led to the recruitment of RomR-GFP at the droplet interface, indicating that this interaction is also detectable in this system **(Fig. 7H)**. Importantly, mixing red-labeled MglB with unlabeled RomR and MglA_GTP488_ led to diffuse localisation of MglA **(Fig. 7I)**. This is due to the formation of the MglB-RomR complex because RomR-GFP was recruited to the lipid interface together with MglB when all three proteins were mixed and MglA was not labeled **(Fig. 7J)**. Thus, the RomR-MglB complex effectively competes with the MglA-MglB complex in oil droplets.

## Discussion

The polarity axis formed by MglA and MglB has long been considered to be formed via an exclusion mechanism linked to the activation of GTP hydrolysis at the lagging cell pole. This mechanism was proposed based on strong evidence: (i) *in vitro* studies on MglA and MglB homologs purified from *T. thermophilus* (MglB_tt_) showed that MglB_tt_ only binds to MglA_GTPtt_ and (ii), *in vivo*, an MglAQ82A/L that cannot hydrolyze GTP, even in the presence of MglB, localizes symmetrically in *M. xanthus* cells (Treuner-Lange *et al*, 2015). Surprisingly, when Baranwal et al. (Baranwal *et al*, 2019) elucidated the structure of the *M. xanthus* MglA-MglB complex, they found that MglB remains associated to MglA_GDP_ after hydrolysis, which is due to the presence of a Ct-helix, a motif that is not present in MglB_tt_. This finding raised profound questions about how cell polarity is established via MglA and MglB and prompted this study. We re-examined how MglA interacts with MglB and how this affects the MglA hydrolysis cycle. We discovered that the MglA system is not controlled by a canonical GAP/GEF system as originally proposed, but is centrally regulated by the RomRX complex, which acts both as an MglA effector and a regulator of the MglA-MglB interaction. We discuss below how these interactions could lead to the spatial partitioning of MglA in the *Myxococcus* cell.

Cell polarity is dictated by a complex interaction network involving MglA, RomRX and MglA effectors at the leading pole, and MglA, MglB, RomRX, RomY and MglC at the lagging cell pole. Carreira et al. (Carreira *et al*, 2023) proposed that cell polarity could be established if interactions between RomR and MglB could be blocked at the leading pole and conversely formed at the lagging pole. Combined with the recent discovery of MglC as a localization factor of MglB, this study provides molecular roots to the model of Carreira et al. and suggests the following scenario:

1. At the leading cell pole, RomRX functions as an MglA effector, recruiting MglA_GTP_. The high concentration of MglA_GTP_ prevents formation of the MglB-MglC complex and thus excludes MglB from the leading cell pole. The interaction of MglA with RomRX is transient and most critical immediately after the polarity switch, as RomRX relocalizes to the lagging pole within a few minutes. During this time, MglA_GTP_ associates with polar effectors such as SgmX for the Type-IV pili and the GltJ ZnR-GYF motif for the Agl-Glt complex. In these complexes, MglA adopts a third state insensitive to the action of MglB, further reinforcing polar localization (Galicia *et al*, 2019) **(Fig. 8)**.
2. At the lagging pole, the spatial localization of GAP activity is essential to establish the polarity axis and the disassembly of the Agl-Glt complex (Attia *et al*, 2024). At this pole, MglB is recruited by MglC, which is favored by the low concentration of MglA (Carreira *et al*, 2023). However, since MglB does not dissociate from MglA_GDP_, a dissociation factor is needed to ensure MglA release following nucleotide hydrolysis. Based on direct *in vitro* evidence, we propose that the RomRX complex is the dissociation factor, acting between reversals when its concentration is high at the lagging pole **(Fig. 8)**. This hypothesis is strongly supported by the co-localization of MglA and MglB at both cell poles observed in a *romR* mutant (Zhang *et al*, 2012b).

**Figure 8:**
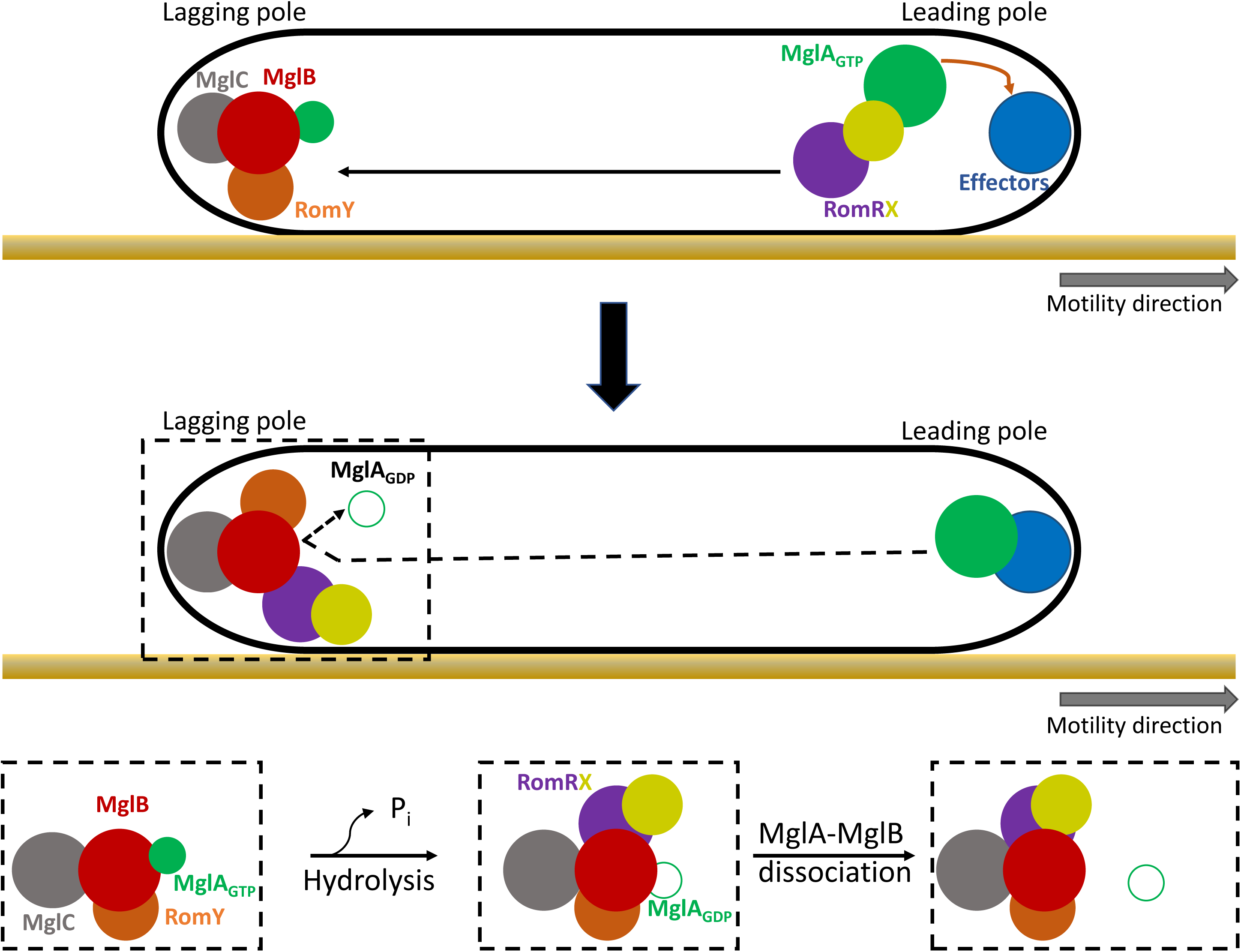
Role of the RomRX complex in establishing cell polarity in M. xanthus. Upper Panel: At the leading pole, the RomRX complex regulates polarity by interacting with MglA_GTP_ through RomX. This interaction supports the recruitment of MglA_GTP_ to the pole where it can associate with polar motility specific effectors present in the A- and S- motility complex (such as SgmX for the Type-IV pili and the GltJ ZnR- GYF motif for the Agl-Glt complex). At the lagging pole, the low concentration of MglA facilitates the recruitment of MglB by MglC. **Middle panel:** RomRX detaches from the leading and it relocalizes to the lagging pole possibly via several interactions (ie MglC, MglB and unknown). As it accumulates at high concentrations, RomRX competes with the formation of the MglA-MglB complex interacting with MglB. **Lower Panel**: However, as observed in vitro, this competition does not entirely block the formation of the MglA-MglB complex due to its low affinity. Given that formation of the MglA-MglB complex and GTP hydrolysis occur rapidly, and that the dissociation of the MglA-MglB complex is the rate-limiting step, RomRX mostly acts as a dissociation factor following hydrolysis.Additionally, RomY plays a crucial role in orienting MglB function toward a GAP in vivo, enhancing its GAP activity and thus accelerating the reaction. This regulation is essential to drive GTP hydrolysis efficiently, as the k_cat_ of MglB alone is low.

Thus, functioning as a polar effector, RomRX first ensures that MglA_GTP_ localizes to the leading pole, while blocking access to the lagging pole by forming a MglB-RomRX complex. Consequently, the common view that the Mgl system is a prokaryotic version of a classical small G-protein system, regulated by spatially localized GAP (MglB) and GEF (RomRX), is obsolete. So how can this system function without a *bone fide* GEF and GAP system? Considering all available data, a view emerges that distinct signaling hubs, one formed by MglA (which is insensitive to the action of MglB (Galicia *et al*, 2019)), the other formed by MglB, segregate at opposite cell poles. The lagging pole system is formed by numerous interactions, between RomR and MglC, MglC and MglB (Carreira *et al*, 2023) and now MglB and RomR. This system is thus the site of numerous regulations: as discussed above it can be destabilized by RomRX and there are likely a number of additional regulators acting on it. There are currently two examples of such regulation:

i. RomY which activates the GAP activity and may be critical to accelerate the reaction *in vivo* because the *k_cat_* of MglB alone is exceedingly low.
ii. Any regulation that modulates interaction of the MglB Ct-helix with MglA converts MglB into a bona fide GAP (Chakraborty *et al*, 2024). Such regulation occurs in the Agl-Glt complex where the GltJ ZnR domain recruits and converts it into a GAP via a direct action on this helix (Attia *et al*, 2024).

Since MglB does not dissociate from MglA, it can neither be called a GAP nor a GEF. Remarkably, the intervention of a dissociation factor could direct the reaction towards either of these functions, depending on how fast the dissociation occurs and whether it competes with or allows GTP hydrolysis (which again is a slow step). Since RomR dissociates both GTP and GDP-bound MglA from MglB it could help direct either reaction depending on the context. In the cell it is clearly the GAP that dominates (possibly due to the presence of RomY, (Szadkowski *et al*, 2022) at the lagging pole between reversals. The notable asymmetry of the complex formed between MglA and the MglB dimer, along with recent findings that the Ct-helix of the proximal MglB protomer is sufficient to stabilize MglA in its GDP- and GTP-bound states (Chakraborty *et al*, 2024), suggests that the second MglB protomer provides an important binding interface for additional regulators (Szadkowski *et al*, 2022). In combination with RomR, these regulators could thus functionalize the complex as a GAP or a GEF.

This emerging picture makes it very difficult to propose a precise reversal mechanism at this stage. We propose that when they become phosphorylated, FrzX and FrzZ each act on the polar signaling complexes to release MglA_GTP_ and allow its relocalization at the lagging pole. The molecular targets of these regulators are yet to be discovered and could involve yet unidentified polar proteins. Nevertheless, this work makes apparent that modulating the partitioning function of RomRX might be a key. Indeed, it can be imagined that promoting the formation of MglA-MglB complexes by alleviating the inhibition of RomRX could trigger a domino effect leading to the release of MglB from MglC and its relocalization at the opposite pole.

Recently, structural homologs of MglA and MglB have been found in *Asgard Archaea*, demonstrating an ancient origin of these systems (Tran et al., 2024). Remarkably, Thor-Rab, the MglA (and Rab) homolog, also contains an arginine residue positioned similarly to the MglA Arg53, suggesting that interaction with the archaeal MglB counterpart, Thor-RB, might lead to a Thor complex functioning with similar enzymatic properties as MglAB. Thus, in all three domains of life, Roadblock domain-containing proteins (as well as the structurally related Longins) may form dimeric platforms that often directly interact with small GTPases (Koonin & Aravind, 2000; Levine *et al*, 2013; Liu *et al*, 2021). In eukaryotic cells, these proteins primarily function as GEFs, but occasionally also as GAPs, through allosteric mechanisms or by incorporating additional regulatory components (Levine et al., 2013; Jansen & Hurley, 2023; Cabrera et al., 2014; Miertzschke et al., 2011). One notable example is the Ragulator complex in the mTOR pathway, where multiple Roadblock domain-containing proteins act as a scaffold for Rag GTPases, again creating a multi-layered regulatory hub crucial for mTOR signaling. As discussed above, the MglB system might also be viewed as such and all these systems could be modular signaling units. In conclusion, studying the MglA-MglB system may reveal universal features of these ubiquitous regulatory modules.

## Methods

### Methods and protocols

#### Protein expression, purification and labeling

##### _-_ Expression and purification of MglA-His_6_, MglAQ82L-His_6_, MglB-His_6_, MglB^ΔCt^-His_6_, SgmX-Cter and RomX-His_6_

The gene sequences of *mglA, m***glAQ82L,** *mglB, mglB^ΔCt^, sgmX-Cter and romX* from *M. xanthus* were inserted in a pET28a plasmid for expression of C-terminal hexa-histidine tagged proteins. Bacterial strains and primers used in this study are listed in Appendix Tables S1 and S2. Plasmid sequences were verified by Sanger sequencing (Eurofins GATC-Biotech, Germany). All constructs were transformed in the BL21(DE3) *E. coli* strain. Cells were grown in Luria-Bertani (LB) medium at 32^°^C and induced at OD_600nm_ 0.5-0.8 for 3 hours by the addition 0.5 mM IPTG (isopropyl-h-d-thiogalactopyranoside). Cells were harvested by centrifugation and pellets were resuspended in buffer A (10 mM Tris-HCl, pH = 7.4, 50 mM NaCl, 5 mM MgCl2, 10 mM Imidazole, pH = 7.4) supplemented with cOmplete™ EDTA-free protease inhibitor cocktail (Roche) and 20 µg/ml DnaseI (Merck). Cells were lysed with a French press at 1 kbar and lysates were centrifuged at 18,000 rpm for 20 minutes at 4^°^C. Supernatants were loaded on 2 ml HisTrap beads columns (GE Healthcare) functionalized with Nickel and equilibrated with Buffer A. After 10 mn of incubation, the resin was washed with 5-column volume of Buffer A and 5-column volume of Buffer B (Buffer A containing 75 mM Imidazole). The protein was then eluted with Buffer C (Buffer A containing 200 mM Imidazole). Fractions containing the protein were pooled, concentrated, dialysed overnight in Buffer D (Buffer A without imidazole) and stored at -80^°^C in Buffer D. Protein purity was analyzed by SDS-PAGE, and protein concentrations were quantified using NanoDrop™ (Thermo Scientific™). The following extinction coefficients, calculated based on the amino acid sequence of the constructed proteins using ProtParam from ExPASy, were used: 16,515 M^-1^·cm^-1^ for MglA-His6 and MglAQ82L-His6, 2,980 M^-1^·cm^-1^ for MglB-His_6_ and MglBΔCt-His_6_, 11,920 M^-1^·cm^-1^ for SgmX-Cter-His_6_, and 8,480 M^-1^·cm^-1^ for RomX-His_6_.

##### _-_ Expression and purification of RomR-His_6_ and RomR_GFP_-His_6_

Synthetic coding sequences of *RomR* and *RomR_GFP_*were obtained from Twist Bioscience, cloned in a modified pET28 vector for expression of C-terminal hexa-histidine tagged proteins. Constructs were transformed in the LEMO21(DE3) *E. coli* strain. Cells were grown overnight in LB medium at 37^°^C and transferred into NZY auto-induction LB medium (NZYtech) for 24 hours at 18^°^C for overexpression. Cells were harvested by centrifugation and pellets were resuspended in Buffer E (50 mM tris-HCl, pH = 8.0, 300 mM NaCl, 5 mM MgCl_2_, 10 mM imidazole, pH = 8.0) supplemented with cOmplete™ EDTA-free protease inhibitor cocktail (Roche) and 20 µg/ml DnaseI. Cells were lysed with a French press at 1 kbar and lysates were centrifuged at 18,000 rpm for 20 minutes at 4^°^C. Supernatants were loaded on a 2 ml HisTrap beads column (GE Healthcare) functionalized with Nickel and equilibrated with a Buffer E. After 10 mn of incubation, the resin was washed with 5-column volume of Buffer E and 5-column volume of Buffer F (Buffer A containing 75 mM Imidazole). The protein was then eluted with Buffer G (Buffer A containing 200 mM Imidazole). Fractions containing the protein were pooled, concentrated, dialyzed overnight in Buffer H (Buffer A without imidazole) for ITC and MST experiments or dialyzed in Buffer D for all other experiments and stored at -80^°^C. Protein purity was analyzed by SDS-PAGE, and protein concentrations were quantified using NanoDrop™ (Thermo Scientific™). The following extinction coefficients were used: 6990 mol^-1^·cm^-1^ for *RomR-His_6_ and*; 26025 mol^-1^·cm^-1^ for *RomR_GFP_-His_6_*.

##### - Labeling of MglA and MglB

Purified MglA_GDP_ and MglB were labeled overnight at 4°C under gentle agitation in the presence of a 5-fold excess of Alexa Fluor™ 488 C5-maleimide (ThermoFisher scientific) or Janelia Fluor® 646 Maleimide (TOCRIS) in Buffer E supplemented with 2 mM TCEP. Unbound fluorophore was removed by passing the reaction mixture through a pre-equilibrated protein desalting column (Zeba Spin Desalting columns from ThermoFisher scientific). Labeled proteins were analyzed on SDS-PAGE gel and imaged using ChemiDoc imaging system (Bio-Rad) to quantify the fraction of labeling. Labeled proteins were stored at -80°C.

#### Fluorescence Anisotropy experiments

##### - Fluorescent nucleotides analogs

Fluorescent nucleotides used in this study were purchased from Jena Bioscience (https://www.jenabioscience.com/), specifically EDA-GTP-ATTO-488 (ref. NU-820-488) and EDA-GDP- ATTO-488 (ref. NU-840-488). When needed, fluorescent nucleotide analogs were diluted in 20 mM HEPES pH 7.5 (Colombo *et al*, 2021).

##### - Fluorescence anisotropy

The binding of fluorescent nucleotides analogs to proteins was monitored by measuring changes in anisotropy using excitation and emission wavelengths of 504 nm and 521 nm, respectively. These experiments were performed using a Safas Xenius XC spectrofluorometer (Safas Monaco) controlled by SP2000 software, version 7.8.13.0 (Colombo *et al*, 2021).

##### - Nucleotides exchange assays

For kinetic experiments, MglA_GDP_, either alone or in combination with MglB, MglB^ΔCt^, RomR, RomX, RomRX or SgmX-Ct, was mixed in a volume of 150 µl in Buffer D (supplemented with 2mM TCEP) at the indicated concentrations. Nucleotide exchange was initiated by adding 0.1 µM of the fluorescent nucleotide analog, and the increase in fluorescence anisotropy was recorded over time until equilibrium was reached. The main reaction controlling the rate in this condition is the dissociation of GDP from MglA (k_off_^GDP^). Subsequently, exchange kinetics were monitored by competing the bound fluorescent nucleotides with an excess of unlabeled nucleotides (100 µM GTP or GDP), resulting in a decrease in fluorescence anisotropy due to the release of the fluorescence nucleotide from MglA. Here, the primary reaction rate is controlled by the dissociation of the fluorescent nucleotide from MglA (k_off_^N1*^).

The reactions involved are described by the following equations:

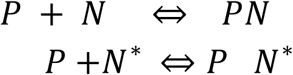

Where P represents MglA_GDP_ or the MglA_GDP_-based protein complex, N is the unlabeled nucleotide, N* is the labeled nucleotide and PN represents the protein-nucleotide complex. These equations implies four rate constants: k_on_^N^, k_off_^N^, k_on_^N*^ and k_off_^N*^.

The binding rate constants (k_on_) cannot be determined by this assay, as they are too fast for anisotropy measurements. However, the dissociation rates corresponding to the slowest reactions can be mathematically described by the following differential equation:

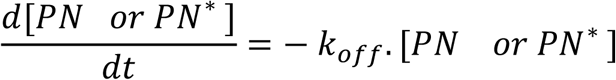

This equation can be resolved by fitting an exponential decay function to estimate the k_off_ value.

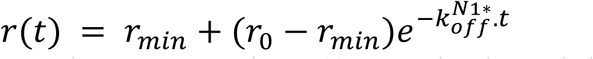

Here, r(t) represents the fluorescence anisotropy at time (t), r_min_ is the minimum anisotropy value for the fluorescent nucleotide alone, r_0_ is the initial fluorescence anisotropy before adding the excess unlabeled nucleotide and k_off_^N*^ is the dissociation rate constants of the fluorescent nucleotide.

- *Steady-state experiment for binding constant determination*

For binding affinity experiments, changes in the fluorescence anisotropy values of EDA-GTP-ATTO-488 (0.1 µM) were measured in the presence of MglA_GDP_ (2 µM) and increasing concentrations of RomX, RomRX (where RomR and RomX were pre-mixed in equal concentrations), or SgmX-Ct. Each measurement was taken after 30 min when steady state was reached. Data were fitted using Igor Pro (version 9.0.5.1) to the binding equation to estimate K_D_:

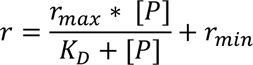

Where r represents the fluorescence anisotropy readout, r_max_ represents the maximum anisotropy value when all the fluorescent nucleotide is bound, [P] is the protein concentration, K_D_ is the binding constant and r_min_ is the minimum anisotropy value corresponding to the fluorescent nucleotide alone in solution. Results were plotted using Igor Pro. Each data point represents the mean of at least three independent measurements, with error bars indicating the standard deviation.

##### - Protein competition experiment

MglA_GDP_ (2 µM) was incubated with MglB (4 µM). The reaction was initiated by adding GTP_488_ or GDP_488_ (0.1 µM) and fluorescence anisotropy was monitored until steady state. Subsequently, RomR, RomX, RomRX or SgmX-ct was added at the indicated concentrations and measurements were continuously taken.

#### GTPase assay

Rates of GTP hydrolysis were measured using the EnzChek Phosphate Assay Kit (E-6646, Thermo Fisher Scientific) based on the method originally described by Webb (Webb, 1992). The assay was performed using a constant concentration of MglA_GDP_ (2 μM) and varying concentration of MglB, MglB^ΔCt^ or RomR-RomX complex, in the presence of 2 mM TCEP, 0.2 mM MESG, PNP (1 U/ml), and a range of GTP concentration as indicated. The reaction was started by the addition of MglA, and phosphate release was recorded by the change in absorbance at 360 nm using 96-well flat-bottom plates (Greiner Bio-One) and a SPARK multimode microplate reader (Tecan) every 30 seconds for at least 30 min. Experiments were repeated a minimum of three times. An R script was used to perform multiple nonlinear regressions on the experimental data to accurately determine the kinetic constants (K_B_, K^’^, K [GDP] and K^’^ [GDP]) by fitting the data to the random or ordered model, and to calculate the coefficient of determination (R²) to assess the goodness of fit.

#### Isothermal Titration calorimetry

Isothermal titration calorimetry (ITC) was employed to determine the thermodynamic parameters and binding constants of the binding interactions between a) MglA_GTPλS or GDP_ and MglB; b) MglA_GTPλS_ and MglB^ΔCt^ and c) RomR or the RomR-RomX complex and MglB .

Case a-b: Experiments were conducted using a MicroCal PEAQ-ITC microcalorimeter (Malvern, UK) at 25°C. Each titration consisted of 19 injections, starting with an initial injection of 0.4 μl followed by 18 injections of 2 μl each. Injections were carried out over 4 seconds with 150-second intervals between injections. The experiments were carried out in Buffer E Supplemented with 5 mM dithiothréitol (DTT). Purified MglA_GDP_ “macromolecule” was placed in the sample cell, while the ligand MglB or MglB^ΔCt^ was loaded into the syringe. Both the cell and the syringe contained 100 μM of GTPλS or GDP. Control experiments involved titrating the ITC buffer into MglA_GDP_ alone to account for heat changes due to dilution. This fitted offset “a constant control heat” was subtracted from the experimental data to accurately calculate the dissociation constant. Data were analyzed using the MicroCal PEAQ-ITC Analysis software, the binding isotherm was fit to a one set of sites model.

Case c: Experiments were performed using a MicroCal PEAQ-ITC microcalorimeter (Malvern, UK) at 12.5°C. Each titration consisted of 13 injections, with the first injection of 0.4 μl and the subsequent 12 injections of 3 μl each. Injections lasted for 4 seconds with 150-second intervals between each injection. The experiments were conducted in Buffer J supplemented with 5 mM DTT. RomR or the RomR-RomX complex (pre-mixed at equal concentrations) was placed in the sample cell, while MglB was loaded into the syringe. A second series of MglB injections with the same parameters was performed, and the resulting ITC data files were concatenated for analysis as a single data file. Control experiments involved titrating the ITC buffer into RomR alone to account for heat changes due to dilution. This fitted offset “a constant control heat” was subtracted from the experimental data to accurately calculate the dissociation constant. Data analysis was performed using the MicroCal PEAQ-ITC Analysis software.

#### Microscale Thermophoresis

Microscale thermophoresis (MST) experiments were conducted using a MonolithX MM-307 instrument (NanoTemper Technologies). MglB was labeled with the Monolith NT Protein Labeling Kit - Red NHS and used at a final concentration of 40 nM. The RomR-RomX complex was titrated in 1:1 serial dilution, ranging from 136 µM to 4 nM. Buffer J supplemented with 5mM DTT and 0.05% tween was used for this experiment. All experiments were performed in standard Monolith Series capillaries (cat# MO-Ko22) with 100% excitation LED power and medium IR-laser power at 24°C. The spectral shift ratio 670nm/650nm was measured as a function of ligand concentration. Each experiment was performed in duplicate to ensure reproducibility. The dissociation constant (K_D_) for the interaction was determined by fitting the thermophoresis data using the MO Affinity Analysis software, assuming a 1:1 binding stoichiometry. Labeled MglB 40nM was bound to RomR with increasing concentrations as shown by MST (Appendix Fig.S5A). The MST F_norm_ values at 650nM with an on time measure at 2.5 seconds in a dose response manner and a signal to noise ratio of 10.3 was used to calculate the binding constant of MglB/RomR . Buffer PBS + 0.05% Tween.

##### Fluorescence microscopy

For each experiment, 1 ml of a CYE-grown culture with an OD of 0.5–1 was centrifuged 5 min at 3000 rpm and resuspended to an OD of 2 in TPM CaCl2 buffer (10 mM Tris-HCl, pH 7.6, 8 mM MgSO_4_, 1 mM KH_2_PO_4_, 1mM CaCl2). Then 2µl of the cell suspension was placed on a glass slide, covered with a 1.5% agar pad, and incubated for 20 min at 32°C before imaging. Time-lapse experiments were performed using an automated inverted epifluorescence Ti2E microscope with Perfect Focus (Nikon), a 100x objective, and a back-illuminated Kinetix Scientific CMOS (sCMOS) camera (Teledyne). All fluorescence images were acquired with minimal exposure time to reduce bleaching and phototoxicity. For time-lapse movies, images were captured every 2 seconds over a 15-minute period.

#### Water-in-oil lipid stabilized emulsion

The lipids used for this experiment included L-α-phosphatidylethanolamine (*E. coli* PE), L-α- phosphatidylglycerol (Egg, Chicken) (sodium salt) (EGG PG), L-α-phosphatidylserine (Brain, Porcine) (sodium salt) (Brain PS), 1’,3’-bis[1,2-dioleoyl-sn-glycero-3-phospho]-glycerol (sodium salt) (cardiolipin 18:1) and L-α-lysophosphatidylcholine (Egg, Chicken) (EGG LysoPC), 1,2-dipalmitoyl-sn-glycero-3- phosphoethanolamine-N-(lissamine rhodamine B sulfonyl) (ammonium salt)(PE*). All lipids were purchased from Merck.

A lipid solution was prepared in a glass vial by mixing 76 mol% *E. coli* PE (or 70 mol% *E.coli* PE + 6 mol% PE* for fluorescent lipids), 4.9 mol% EGG PG, 9.3 mol% cardiolipin 18:1, 6.5mol% PS and 3 mol% Egg Lyso PC in chloroform. This composition was chosen to mimic the inner membrane of *M. xanthus.* The mixture was dried overnight in a vacuum desiccator and then rehydrated in Buffer E to a final concentration of 2mg/ml (≈ 0.83 mM). The resulting suspension was sonicated using a tip sonicator with refrigeration, followed by centrifugation at 1000 rpm for 10 mn to remove titanium particles. The suspensions were stored at 4 °C under nitrogen and used within a week.

To prepare water-in-oil emulsions, the lipid suspension was diluted to a final concentration of 3 µM with the proteins of interest in Buffer E. The oil phase was prepared by dissolving 1 mg/ml cithrol (CRODA) surfactant in Squalene oil (Mreck). To form the emulsion, the water phase was added to the oil phase at a typical volume ratio of 1:20 in a microcentrifuge tube. The mixture was gently mixed by pipetting up and down, avoiding vigorous agitation to maintain emulsion stability. Brief vortex mixing at low speed was used for additional mixing. The successful formation of the droplets was indicated by the uniform cloudiness of the sample. The emulsion was placed in a glass microscope chamber, with droplets positioned between a microscope coverslip and glass slide spaced with double-sided adhesive tape **<u>(Appendix Fig.S4A)</u>**. Droplets and fluorescent proteins were imaged using a Zeiss Axio Observer Z1 microscope equipped with a 100x/1.4NA Oil Ph3 Plan-Apochromat objective and a Hamamatsu ORCA-Flash 4.0LT camera. Images were acquired with Zen 2.3 blue edition.

## Acknowledgements

We acknowledge Jessica Colombo for her help in setting the lipid droplet system. This work was funded by the Centre National de la Recherche Scientifique (CNRS), Aix-Marseille University, the European Research Council to TM (JAWS-Advanced Grant ERC-2019-ADG: 885145). CD was funded by a CENTURI postdoctoral fellowship and by an ERC advanced grant to TM.

## Author contributions

CD, AM, JH and TM designed research. CD performed the experiments. CD and DB-K performed and analyzed ITC experiments, SL performed and analyzed the in vivo data. RV and FD did the first purification of RomR and provided the protocol. CS-K and CD analyzed kinetic experiments and wrote the enzymology model. JH assisted with the initial setup of experiments and contributed valuable insights for interpreting the results. CD, AM and TM analyzed the data, interpreted the results, and wrote the manuscript. TM acquired funding.

## Data Availability Section

All data porting this study are included in the main text, EV figures and Appendix.

## Expanded view figure legends

**Figure EV1:**
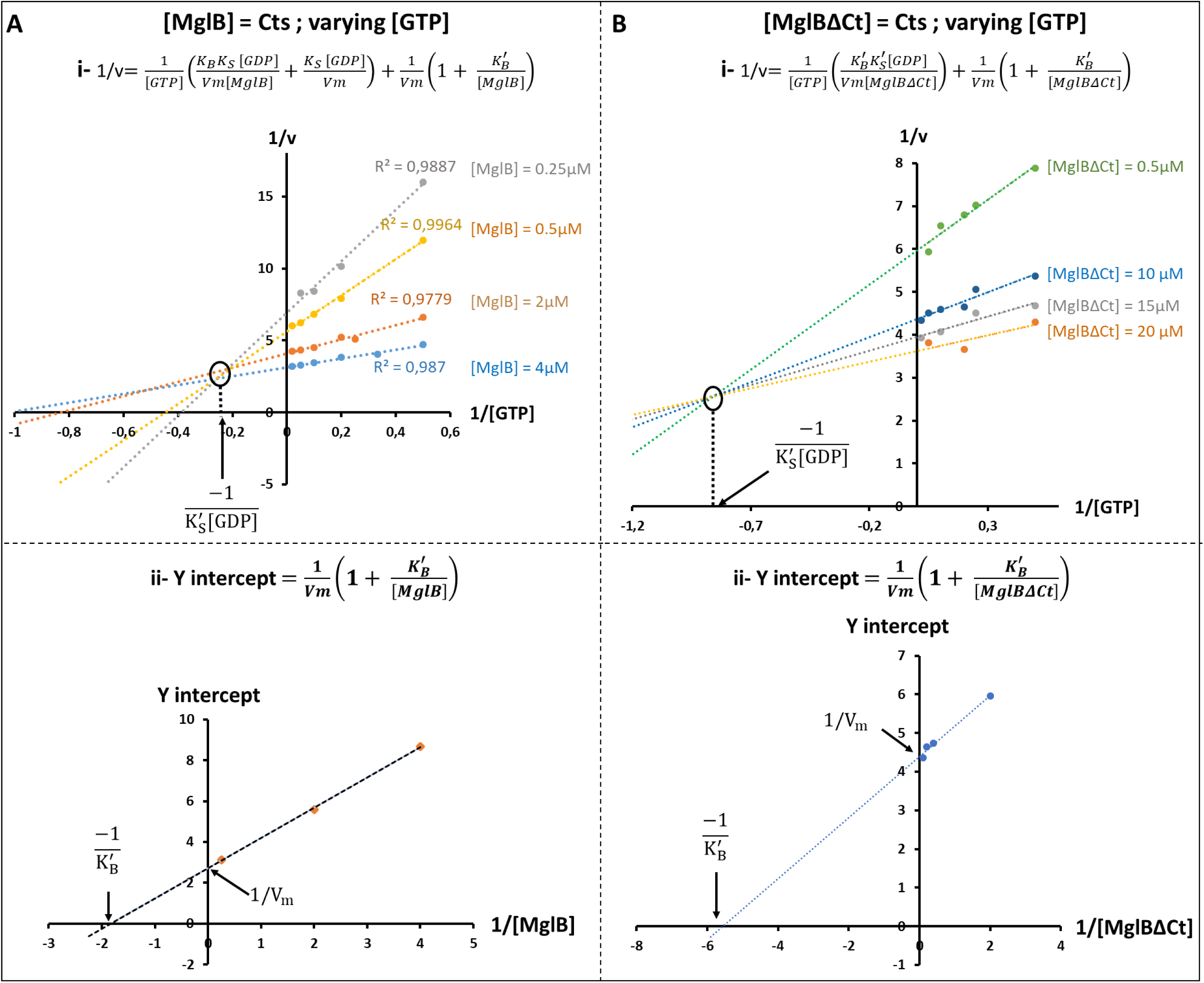
Kinetic analysis of MglA toward GTP and MglB or MglB^ΔCt^. **A. Top:** Plot of 1/v as a function of 1/[GTP] at varying MglB concentrations (0.25, 0.5, 2 and 4µM). The data were modeled using the equation (i). All lines converge to a single point corresponding to -1/K_s_^’^ [GDP], highlighting the substrate-dependent modulation of enzymatic activity. **Bottom:** Following equation (ii), the y-intercept (corresponding to the plot of 1/v in the function of 1/[GTP] at a single MglB concentration) was plotted in the function of 1/[MglB] at three different MglB concentrations 0.25, 0.5 and 4 µM. Y-intercept corresponds to 1/Vm and the x-intercept corresponds to -1/K ^’^. **B. Top:** Plot of 1/v as a function of 1/[GTP] at varying MglB^ΔCt^ concentrations (0.5, 10, 15 and 20µM). The data were modeled using the equation (i). All lines converge to a single point corresponding to -1/K_s_^’^ [GDP]. **Bottom:** Following equation (ii), the y-intercept (corresponding to the plot of 1/v in the function of 1/[GTP] at a single MglB^ΔCt^concentration) was plotted in the function of 1/[MglB^ΔCt^] at four different MglB^ΔCt^ concentrations 0.5, 2.5, 5 and 10 µM. Y-intercept corresponds to 1/Vm and the x-intercept corresponds to - 1/K ^’^.

## References

1. Allam AH, Charnley M & Russell SM (2018) Context-Specific Mechanisms of Cell Polarity Regulation. Journal of Molecular Biology 430: 3457–3471

2. Attia B, My L, Castaing JP, Dinet C, Le Guenno H, Schmidt V, Espinosa L, Anantharaman V, Aravind L, Sebban-Kreuzer C, et al (2024) A molecular switch controls assembly of bacterial focal adhesions. Science Advances 10: eadn2789

3. Bagshaw C (2001) ATP analogues at a glance. J Cell Sci 114: 459–460

4. Baranwal J, Lhospice S, Kanade M, Chakraborty S, Gade PR, Harne S, Herrou J, Mignot T & Gayathri P (2019) Allosteric regulation of a prokaryotic small Ras-like GTPase contributes to cell polarity oscillations in bacterial motility. PLoS Biol 17: e3000459

5. Bautista S, Schmidt V, Guiseppi A, Mauriello EMF, Attia B, Elantak L, Mignot T & Mercier R (2023) FrzS acts as a polar beacon to recruit SgmX, a central activator of type IV pili during Myxococcus xanthus motility. The EMBO Journal 42: e111661

6. Bos JL, Rehmann H & Wittinghofer A (2007) GEFs and GAPs: Critical Elements in the Control of Small G Proteins. Cell 129: 865–877

7. Carreira LAM, Szadkowski D, Lometto S, Hochberg GKA & Søgaard-Andersen L (2023) Molecular basis and design principles of switchable front-rear polarity and directional migration in Myxococcus xanthus. Nat Commun 14: 4056

8. Carreira LAM, Szadkowski D, Müller F & Søgaard-Andersen L (2022) Spatiotemporal regulation of switching front–rear cell polarity. Current Opinion in Cell Biology 76: 102076

9. Carreira LAM, Tostevin F, Gerland U & Søgaard-Andersen L (2020) Protein-protein interaction network controlling establishment and maintenance of switchable cell polarity. PLOS Genetics 16: e1008877

10. Chakraborty S & Gayathri P (2024) RomX, a novel prokaryotic regulator, links the response receiver domain of RomR with GTP-bound MglA for establishing *Myxococcus xanthus* polarity. doi:10.1101/2024.02.20.581209 [PREPRINT]

11. Chakraborty S, Kanade M & Gayathri P (2024) Mechanism of GTPase activation of a prokaryotic small Ras-like GTPase MglA by an asymmetrically interacting MglB dimer. Journal of Biological Chemistry 300: 107197

12. Cherfils J & Zeghouf M (2013) Regulation of Small GTPases by GEFs, GAPs, and GDIs. Physiological Reviews 93: 269–309

13. Chiou J-G, Balasubramanian MK & Lew DJ (2017) Cell Polarity in Yeast. Annu Rev Cell Dev Biol 33: 77–101

14. Colombo J, Antkowiak A, Kogan K, Kotila T, Elliott J, Guillotin A, Lappalainen P & Michelot A (2021) A functional family of fluorescent nucleotide analogues to investigate actin dynamics and energetics. Nat Commun 12: 548

15. Dinet C & Mignot T (2023) Unorthodox regulation of the MglA Ras-like GTPase controlling polarity in Myxococcus xanthus. FEBS Lett 597: 850–864

16. Faure LM, Fiche J-B, Espinosa L, Ducret A, Anantharaman V, Luciano J, Lhospice S, Islam ST, Tréguier J, Sotes M, et al (2016) The mechanism of force transmission at bacterial focal adhesion complexes. Nature 539: 530–535

17. Galicia C, Lhospice S, Varela PF, Trapani S, Zhang W, Navaza J, Herrou J, Mignot T & Cherfils J (2019) MglA functions as a three-state GTPase to control movement reversals of Myxococcus xanthus. Nat Commun 10: 5300

18. Goryachev AB & Leda M (2019) Autoactivation of small GTPases by the GEF–effector positive feedback modules. F1000Res 8: F1000 Faculty Rev-1676

19. Guzzo M, Murray SM, Martineau E, Lhospice S, Baronian G, My L, Zhang Y, Espinosa L, Vincentelli R, Bratton BP, et al (2018) A gated relaxation oscillator mediated by FrzX controls morphogenetic movements in Myxococcus xanthus. Nat Microbiol 3: 948–959

20. Hennig A, Markwart R, Esparza-Franco MA, Ladds G & Rubio I (2015) Ras activation revisited: role of GEF and GAP systems. Biological Chemistry 396: 831–848

21. Herrou J & Mignot T (2020) Dynamic polarity control by a tunable protein oscillator in bacteria. Current Opinion in Cell Biology 62: 54–60

22. Houston DW (2017) Cell Polarity in Development and Disease

23. Iden S & Collard JG (2008) Crosstalk between small GTPases and polarity proteins in cell polarization. Nat Rev Mol Cell Biol 9: 846–859

24. Ivanetich KM & Goold RD (1989) A rapid equilibrium random sequential bi-bi mechanism for human placental glutathione S-transferase. Biochimica et Biophysica Acta (BBA) - Protein Structure and Molecular Enzymology 998: 7–13

25. Kanade M, Singh NB, Lagad S, Baranwal J & Gayathri P (2021) Dual specificity of a prokaryotic GTPase-activating protein (GAP) to two small Ras-like GTPases in Myxococcus xanthus. FEBS J 288: 1565–1585

26. Keilberg D, Wuichet K, Drescher F & Søgaard-Andersen L (2012) A Response Regulator Interfaces between the Frz Chemosensory System and the MglA/MglB GTPase/GAP Module to Regulate Polarity in Myxococcus xanthus. PLOS Genetics 8: e1002951

27. Kirkpatrick CL & Viollier PH (2011) Poles Apart: Prokaryotic Polar Organelles and Their Spatial Regulation. Cold Spring Harb Perspect Biol 3: a006809

28. Koonin EV & Aravind L (2000) Dynein light chains of the Roadblock/LC7 group belong to an ancient protein superfamily implicated in NTPase regulation. Curr Biol 10: R774–776

29. Kühn MJ, Macmillan H, Talà L, Inclan Y, Patino R, Pierrat X, Al-Mayyah Z, Engel JN & Persat A (2023) Two antagonistic response regulators control Pseudomonas aeruginosa polarization during mechanotaxis. The EMBO Journal 42: e112165

30. Laloux G & Jacobs-Wagner C (2014) How do bacteria localize proteins to the cell pole? J Cell Sci 127: 11–19

31. Leonardy S, Miertzschke M, Bulyha I, Sperling E, Wittinghofer A & Søgaard-Andersen L (2010) Regulation of dynamic polarity switching in bacteria by a Ras-like G-protein and its cognate GAP. The EMBO Journal 29: 2276–2289

32. Levine TP, Daniels RD, Wong LH, Gatta AT, Gerondopoulos A & Barr FA (2013) Discovery of new Longin and Roadblock domains that form platforms for small GTPases in Ragulator and TRAPP-II. Small GTPases 4: 62–69

33. Liu Y, Makarova KS, Huang W-C, Wolf YI, Nikolskaya AN, Zhang X, Cai M, Zhang C-J, Xu W, Luo Z, et al (2021) Expanded diversity of Asgard archaea and their relationships with eukaryotes. Nature 593: 553–557

34. MacCready JS, Hakim P, Young EJ, Hu L, Liu J, Osteryoung KW, Vecchiarelli AG & Ducat DC (2018) Protein gradients on the nucleoid position the carbon-fixing organelles of cyanobacteria. Elife 7: e39723

35. Mauriello E (2019) How bacteria arrange their organelles. Elife 8: e43777

36. Mayor R & Etienne-Manneville S (2016) The front and rear of collective cell migration. Nat Rev Mol Cell Biol 17: 97–109

37. Mercier R, Bautista S, Delannoy M, Gibert M, Guiseppi A, Herrou J, Mauriello EMF & Mignot T (2020) The polar Ras-like GTPase MglA activates type IV pilus via SgmX to enable twitching motility in Myxococcus xanthus. Proceedings of the National Academy of Sciences 117: 28366–28373

38. Mercier R & Mignot T (2016) Regulations governing the multicellular lifestyle of *Myxococcus xanthus*. Current Opinion in Microbiology 34: 104–110

39. Miao W, Eichelberger L, Baker L & Marshall MS (1996) p120 Ras GTPase-activating Protein Interacts with Ras-GTP through Specific Conserved Residues. Journal of Biological Chemistry 271:15322–15329

40. Miertzschke M, Koerner C, Vetter IR, Keilberg D, Hot E, Leonardy S, Søgaard-Andersen L & Wittinghofer A (2011) Structural analysis of the Ras-like G protein MglA and its cognate GAP MglB and implications for bacterial polarity. The EMBO Journal 30: 4185–4197

41. Mignot T, Shaevitz JW, Hartzell PL & Zusman DR (2007) Evidence that focal adhesion complexes power bacterial gliding motility. Science 315: 853–856

42. Piroli ME, Blanchette JO & Jabbarzadeh E (2019) Polarity as a physiological modulator of cell function. Front Biosci (Landmark Ed) 24: 451–462

43. Posner I, Engel M & Levitzki A (1992) Kinetic model of the epidermal growth factor (EGF) receptor tyrosine kinase and a possible mechanism of its activation by EGF. Journal of Biological Chemistry 267: 20638–20647

44. Rowlett VW & Margolin W (2013) The bacterial Min system. Curr Biol 23: R553–556

45. Schumacher D & Søgaard-Andersen L (2017) Regulation of Cell Polarity in Motility and Cell Division in Myxococcus xanthus. Annu Rev Microbiol 71: 61–78

46. Szadkowski D, Carreira LAM & Søgaard-Andersen L (2022) A bipartite, low-affinity roadblock domain-containing GAP complex regulates bacterial front-rear polarity. PLoS Genet 18: e1010384

47. Szadkowski D, Harms A, Carreira LAM, Wigbers M, Potapova A, Wuichet K, Keilberg D, Gerland U & Søgaard-Andersen L (2019) Spatial control of the GTPase MglA by localized RomR–RomX GEF and MglB GAP activities enables Myxococcus xanthus motility. Nat Microbiol 4: 1344–1355

48. Treuner-Lange A, Macia E, Guzzo M, Hot E, Faure LM, Jakobczak B, Espinosa L, Alcor D, Ducret A, Keilberg D, et al (2015) The small G-protein MglA connects to the MreB actin cytoskeleton at bacterial focal adhesions. J Cell Biol 210: 243–256

49. Webb MR (1992) A continuous spectrophotometric assay for inorganic phosphate and for measuring phosphate release kinetics in biological systems. Proceedings of the National Academy of Sciences 89: 4884–4887

50. Wettmann L & Kruse K (2018) The Min-protein oscillations in Escherichia coli: an example of self-organized cellular protein waves. Philos Trans R Soc Lond B Biol Sci 373: 20170111

51. Wittinghofer A, Scheffzek K & Ahmadian MR (1997) The interaction of Ras with GTPase-activating proteins. FEBS Letters 410: 63–67

52. Zhang Y, Ducret A, Shaevitz J & Mignot T (2012a) From individual cell motility to collective behaviors: insights from a prokaryote, Myxococcus xanthus. FEMS Microbiol Rev 36: 149–164

53. Zhang Y, Franco M, Ducret A & Mignot T (2010) A Bacterial Ras-Like Small GTP-Binding Protein and Its Cognate GAP Establish a Dynamic Spatial Polarity Axis to Control Directed Motility. PLoS Biol 8: e1000430

54. Zhang Y, Guzzo M, Ducret A, Li Y-Z & Mignot T (2012b) A Dynamic Response Regulator Protein Modulates G-Protein–Dependent Polarity in the Bacterium Myxococcus xanthus. PLOS Genetics 8: e1002872

